# Radioligand therapy in combination with CAR T cells overcomes the heterogeneous immunosuppressive prostate tumor microenvironment

**DOI:** 10.64898/2026.07.02.736191

**Authors:** Jiangyue Liu, Iveta Fajnorova, Yuwei Ren, Kofi Poku, Shuai Yang, Yu-Hsuan Fu, Cari A. Young, Lupita S. Lopez, Reginaldo C. A. Rosa, Handan Hong, Jingting Hao, Deyi Chen, Pauline Jeanjean, Ines Camille Azrour, Ava Fakharpour, Lea Christian, John P. Murad, Yukiko Yamaguchi, Laura H. Porter, Vikram Adhikarla, Russell Rockne, Stephen J. Forman, Yun Rose Li, Tanya B. Dorff, Gail R. Risbridger, Renea Taylor, Christine E. Mona, Saul J. Priceman

## Abstract

^177^Lu-PSMA-617 (Pluvicto^TM^, Lu-177 RLT) is an FDA-approved targeted radioligand therapy (RLT) for metastatic castration-resistant prostate cancer (mCRPC), but its durability of response to this singular approach poses a challenge to the field. Chimeric antigen receptor (CAR) T cell therapy has revolutionized clinical practice for hematological malignancies, but its clinical development for solid tumors, including mCRPC, has been encumbered by antigen heterogeneity and the immunosuppressive tumor microenvironment (TME). Here, we evaluate the therapeutic combination of Lu-177 RLT and PSCA-CAR T cells to overcome these barriers. In human xenograft and mouse syngeneic prostate cancer models with homogeneous or heterogeneous antigen expression, the sequential administration of Lu-177 RLT, cyclophosphamide (Cy), and PSCA-CAR T cells improves tumor control and prolongs survival compared to monotherapies. Mechanistically, Lu-177 RLT alone or with Cy remodels the TME by promoting pro-inflammatory myeloid responses and activating endogenous T cells, while enhancing CAR T cell activation and effector function. We additionally evaluated ^225^Ac-PSMA-617 RLT as an emerging approach in combination with CAR T cells and observed anti-tumor responses, supporting its potential as an alternative RLT partner. These findings support RLT as an immune priming strategy to enhance CAR T cell therapy and provide a rationale for clinical translation of this combination in mCRPC.

**One Sentence Summary:** Combining ^177^Lu-PSMA-617 radioligand therapy with PSCA-CAR T cells improves tumor control and survival in prostate cancer models by overcoming the antigen heterogeneity and reshaping the immunosuppressive tumor microenvironment.

## INTRODUCTION

Metastatic castration-resistant prostate cancer (mCRPC) remains a lethal disease, with a median survival of 25.6 months (*1*). Despite recent advances, no therapy can reliably eliminate mCRPC tumors or prevent relapse. Recurrences are largely driven by multiple treatment-resistance mechanisms or emergence/selection of therapy-resistant clones, highlighting the need for combinatorial strategies that can overcome resistance to therapy and provide long-term durability in response. ^177^Lu-PSMA-617 (Pluvicto^TM^, Lu-177) is a recent FDA-approved radioligand therapy (RLT) that represents a major clinical advance for patients with mCRPC, selectively targeting the highly expressed prostate-specific membrane antigen (PSMA) to deliver tumor-directed radiation (*2*). While Lu-177 RLT improves survival in patients, responses are rarely durable (*3, 4*). Resistance is thought to arise from tumor antigen heterogeneity and intrinsic or acquired radioresistance, thereby limiting long-term efficacy.

Chimeric antigen receptor (CAR) T cell therapy is a cellular immunotherapy that engineers patient T cells to target tumor-associated cell surface antigens. Several CAR targets are being clinically evaluated for the treatment of mCRPC (*5–7*). Our group has demonstrated safety and promising responses of CAR T cells targeting prostate stem cell antigen (PSCA) for mCRPC (*8*). However, responses are constrained by antigen heterogeneity and a profoundly immunosuppressive tumor microenvironment (TME), highlighting the need for novel combinatorial and/or T cell engineering approaches that simultaneously address both heterogeneity and immune suppression (*9*).

Among the numerous potential combination strategies, radiotherapy and immunotherapy have emerged as ideal combinatorial candidates for the treatment of mCRPC. For instance, recent studies suggest that Lu-177 RLT can induce immunomodulation and sensitize tumors to immune checkpoint blockade (ICB), which is being investigated clinically (*10–12*). In parallel, evidence has shown that focal radiation therapy (RT) shifts the immune landscape by increasing antigen presentation and enhancing CAR T cell trafficking, infiltration, and active functionality (*13–15*). Our group has an ongoing phase 1b trial evaluating metastasis-directed radiotherapy (MDRT) in combination with PSCA-CAR T cells for the treatment of mCRPC (NCT05805371).

Building on these insights, here we preclinically evaluate the efficacy and safety of combining Lu-177 RLT with PSCA-CAR T cell therapy in human xenograft and mouse syngeneic prostate cancer models. We perform a comprehensive assessment of immunological changes upon single or combination treatment, integrating flow cytometry and single-cell RNA sequencing. Our data demonstrate that sequential administration of Lu-177 RLT followed by PSCA-CAR T cell therapy enhances survival and anti-tumor responses in preclinical models. Mechanistically, the combination simultaneously targets antigen heterogeneity and remodels the TME to support sustained anti-tumor activity. We further extend these findings ^225^Ac-PSMA-617 (Ac-225) RLT, supporting the broad potential of combining alpha-emitting RLT and CAR T cell therapy. By applying these two therapeutic modalities that are either FDA approved or under active clinical investigation, we aim to rapidly clinically translate this approach to patients with mCRPC.

## RESULTS

### PSMA and PSCA are heterogeneously expressed in prostate cancer

We first evaluated expression of PSMA and PSCA using clinically relevant models and publicly available datasets. Immunohistochemistry and bulk RNA-sequencing (RNA-seq) analysis of the MURAL collection of prostate cancer patient-derived xenografts (PDXs) (*16*), which includes both primary and metastatic tumors representing diverse pathological states, revealed variability in PSMA and PSCA expression both across different PDX models and within individual tumor sections (**Figure 1a-c**). To extend the findings, we incorporated the Human Prostate Single Cell Atlas (HuPSA), which compiles 74 samples from public datasets with cell type annotations (*17*). Co-expression of PSMA and PSCA was observed in certain cell types (prostate adenocarcinoma, AdPCa), whereas other cell types exhibited heterogeneous expression patterns with limited overlap between PSMA and PSCA (**Figure 1d-e**). Such intra- and inter-tumor antigen heterogeneity highlighted a key limitation of single-antigen targeted therapies, namely, tumor relapse driven by clonal escape of antigen-negative or antigen-low populations. Notably, although PSMA and PSCA are commonly associated with adenocarcinoma, their expression was also detected in mixed neuroendocrine/adenocarcinoma phenotypes (**Figure 1a**). Collectively, these data reveal substantial antigen heterogeneity in prostate cancer, providing a strong rationale for PSMA and PSCA dual-targeting therapeutic strategies.

**Figure 1.**
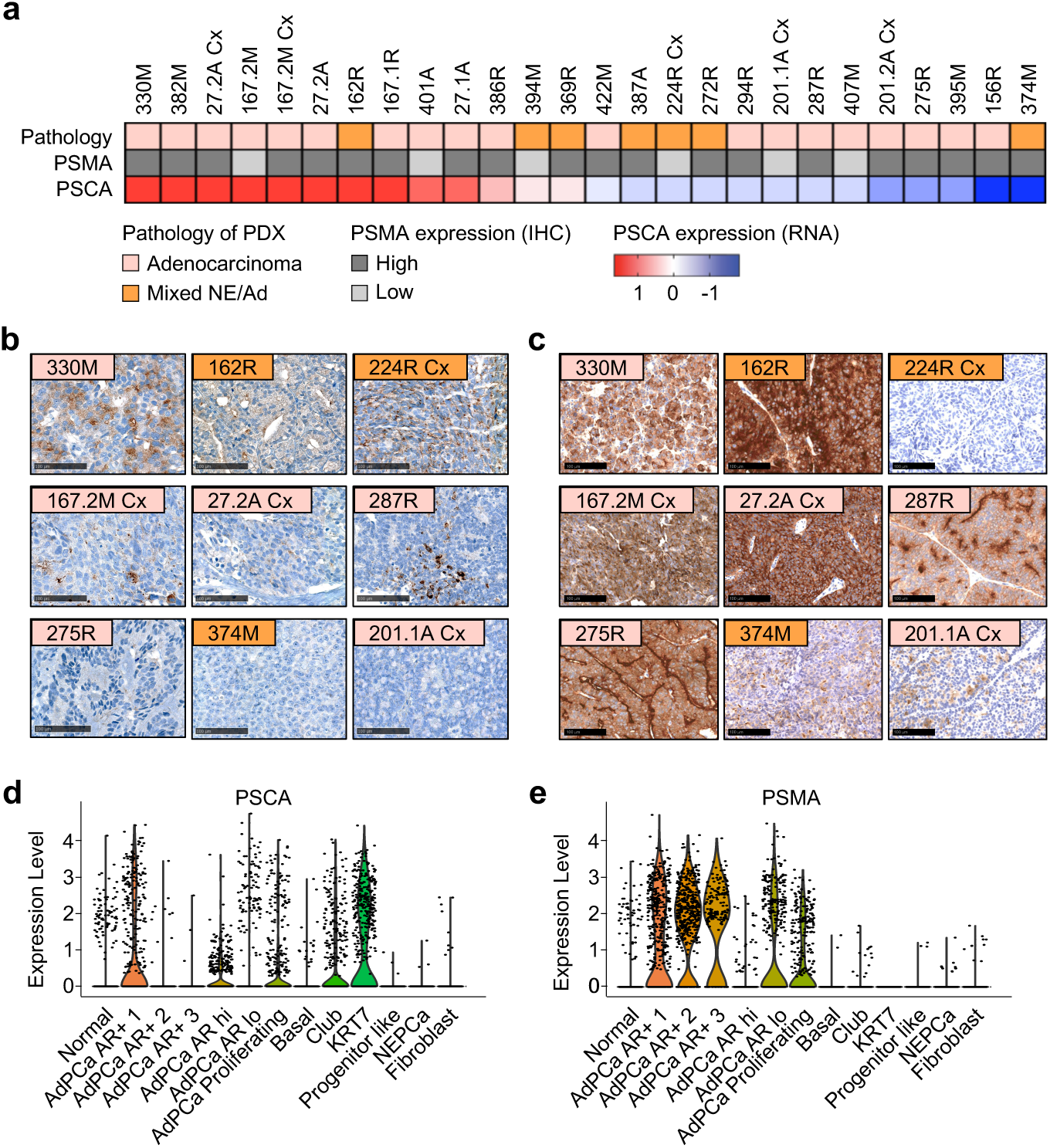
Prostate cancer exhibits antigen heterogeneity across PDX models and public databases. (**a**) Summary of PSMA and PSCA expression in prostate cancer PDX from the MURAL collection. Cx, sublines in castrated mice. (**b-c**) Representative IHC staining of PSCA (**b**) and PSMA (**c**) in selected PDX models, organized from top to bottom based on PSCA expression levels (high, low, and negative). Scale bar, 100 µm. (**d-e**) Transcriptomic expression levels of *FOLH1* (PSMA) and *PSCA* from public datasets (HuPSA).

### Lu-177 RLT and CAR T cell combination enhances survival in prostate cancer xenograft models

To evaluate the therapeutic potential of combining PSMA-targeted RLT with PSCA-targeted CAR T cells, we engineered the human prostate cancer cell line PC3-PIP (PSMA-positive) to express human PSCA and firefly luciferase (ffLuc). Following intracardiac injection to model metastatic disease, these cells established metastases in the liver and bone of NSG mice, which were detected by ^68^Ga-PSMA-11 positron emission tomography/computed tomography (PET/CT) and bioluminescence imaging at day 45 post-injection (**Figure 2a-b**). Lu-177 RLT monotherapy, administered at either 15 or 30 MBq, resulted in modest anti-tumor activity. PSCA-CAR T cells alone significantly prolonged survival compared to untreated (UT) controls (P < 0.0001), but did not produce complete responses (**Figure 2c-d**). In contrast, sequential administration of RLT followed by PSCA-CAR T cells markedly enhanced therapeutic efficacy, leading to tumor regression and long-term survival in 40% of mice. The combination therapy was well-tolerated, with no significant changes observed in body weight (**Figure 2e**). To further assess the hematologic safety, complete blood counts were measured at 3, 7, and 27 days following RLT administration (18, 25, and 42 days post tumor injection). Overall, no significant reductions were observed in white blood cell (WBC), lymphocyte, red blood cell (RBC), or platelet (PLT) at any timepoint compared to the UT controls. A transient increase in PLT was observed at day 42 but was not associated with hematologic toxicity (**Figure 2f-i**).

**Figure 2.**
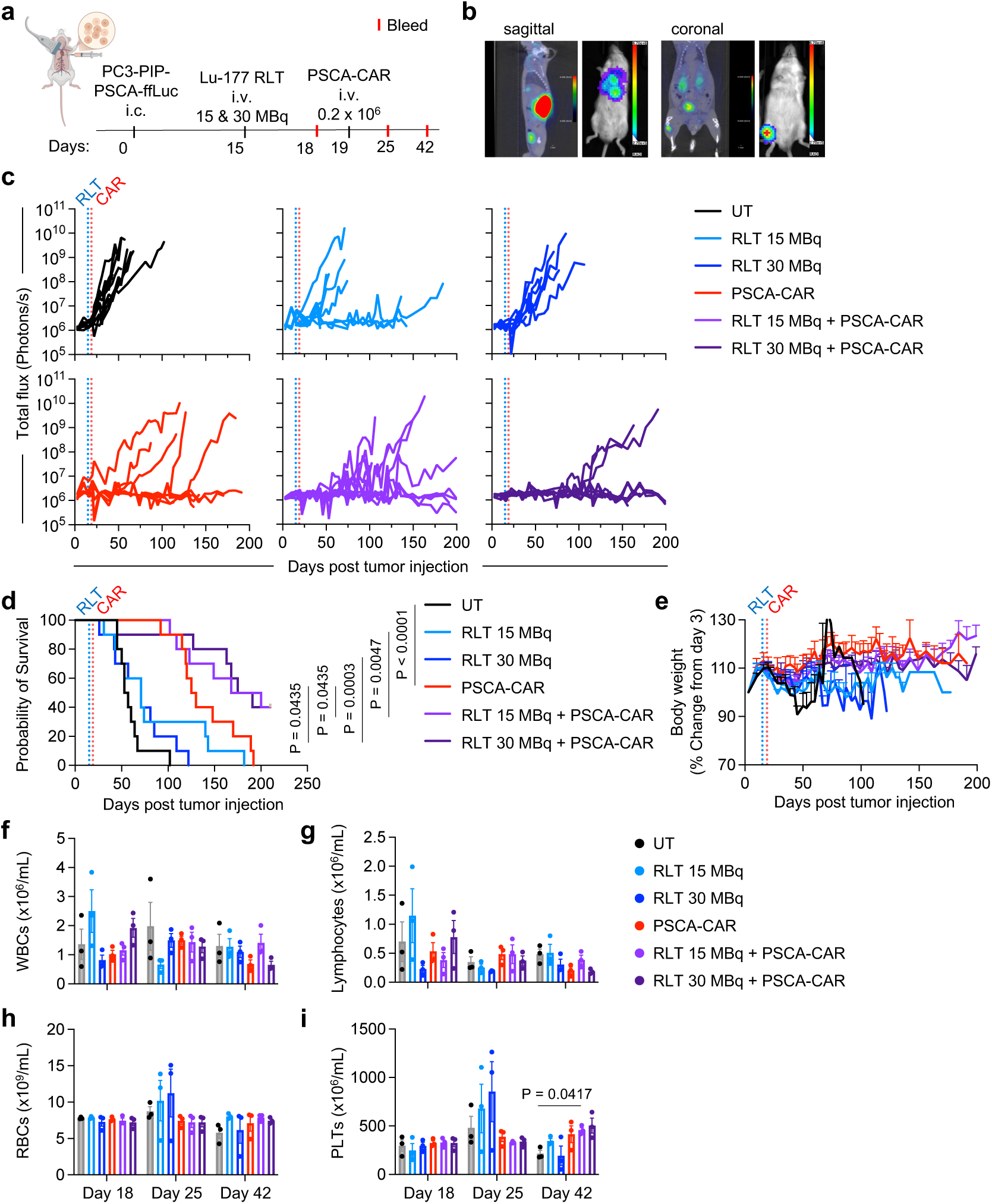
Lu-177 RLT and PSCA-CAR T cell combination enhances therapeutic efficacy and animal survival in a human xenograft prostate cancer model. (**a**) Schematic of tumor injection and treatment schedule in NSG mice bearing metastatic prostate cancer. i.c., intracardiac; i.v., intravenous. On day 15 post-tumor injection, mice received Lu-177 RLT at 15 MBq or 30 MBq (i.v.); Four days later (day 19), mice were treated with 0.2 × 10^6^ PSCA-CAR T cells (i.v.), either as monotherapy or in combination. Blood samples (50 µL) were collected on days 18, 25, and 42 (corresponding to days 3, 7, and 21 post-RLT) for complete blood count (CBC) analysis. (**b**) Representative PET/CT scans (left) and bioluminescence imaging (BLI, right) of untreated mice 45 days post-tumor injection. UT, untreated. (**c**) Longitudinal whole-body flux imaging of PC3-PIP-PSCA-ffLuc metastatic tumor-bearing mice treated with RLT alone, PSCA-CAR T cells alone, or the combination (n = 10 per group). Flux imaging was performed twice a week until day 200, and signals were quantified using a consistent region of interest (ROI) for each animal. (**d**) Kaplan-Meier survival curves for each treatment group. P-value is calculated using multiple comparisons with Holm-Šídák’s correction. (**e**) Body weight changes over time in each group, normalized to body weight at day 3 post-tumor injection. Data are presented as mean ± SEM. (**f-i**) Hematologic analysis by CBC, including white blood cells (WBCs, **f**), lymphocytes (**g**), red blood cells (RBCs, **h**), and platelets (PLTs, **i**) from each group (n = 3 per group). Data are presented as mean ± SEM. Statistical significance was assessed using two-way ANOVA with Geisser-Greenhouse correction, followed by Tukey’s multiple comparison test.

RLT localized to the tumor site may not only irradiate PSMA-expressing cancer cells but also potentially impair the viability of subsequently administered CAR T cells. Radioisotope activity within the tumor decreases over time due to radioactive decay, which may influence the outcome of the combination therapy. To determine whether extending the interval between RLT and CAR T cell administration affects therapeutic outcomes, we compared treatment schedules with a 4-day versus 6-day interval between RLT and CAR T cell treatment (**Figure S1a**). We found that delaying CAR T cell treatment did not significantly alter the therapeutic efficacy or overall survival (**Figure S1b-c**). The extended interval did not result in significant differences in treatment-associated toxicities (**Figure S1d-h**). Collectively, these results demonstrate that RLT priming enhances CAR T cell efficacy in metastatic prostate cancer, prompting us to further investigate the underlying immunological mechanisms of RLT-mediated priming in an immunocompetent setting.

### Lu-177 RLT induces immunomodulatory effects in mouse syngeneic prostate cancer models

While the xenograft models demonstrate the therapeutic benefit of combining RLT with PSCA-CAR T cells, they do not capture the potential contribution of an intact immune system to the combination therapy. To investigate the immunomodulatory effects of Lu-177 RLT within the TME, we employed an immunocompetent syngeneic prostate cancer model. To establish the model, we engineered the murine prostate cancer cell line RM-9 to express human PSMA (hPSMA, RM9-PSMA-ffLuc), human PSCA (hPSCA, RM9-PSCA-ffLuc), or both antigens (RM9-PSMA-PSCA-ffLuc) (**Figure S2a-b**). Lu-177 RLT binding was confirmed *in vitro*, showing antigen-specific association with RM9-PSMA-ffLuc and RM9-PSMA-PSCA-ffLuc cells, with minimal binding to RM9-WT and RM9-PSCA-ffLuc (**Figure S2c**). *In vivo* targeting was further validated by biodistribution analysis. RM9-PSMA-ffLuc cells were engrafted subcutaneously in C57BL/6 mice, followed by administration of a non-therapeutic dose of Lu-177 RLT (**Figure S2d**). Kidney uptake was transient and declined by 6 hours, consistent with the renal clearance, whereas tumor uptake increased by 0.5 hours and remained detectable up to 168 hours (**Figure S2e**). The estimated tumor absorbed dose was 129 mGy/MBq (**Figure S2f** & **Table S1**).

We next evaluated the immunomodulatory effects of Lu-177 RLT in prostate cancer-bearing immunocompetent mice. RM9-PSMA-ffLuc tumor-bearing C57BL/6 mice were treated with Lu-177 RLT at 37, 74, or 120 MBq, and tumors were harvested 6 days post-treatment (**Figure S3a**). Dose selection was based on prior studies (*18*). Flow cytometry analysis revealed dose-dependent remodeling of the myeloid compartment, which constitutes the main immune population in the prostate TME (**Figure S3b-c**). Specifically, the Ly6G^+^ Ly6C^mid^ granulocytic population increased, whereas the Ly6G^-^ Ly6C^-^ tumor-associated macrophage (TAM)-like population decreased, and the Ly6G^-^ Ly6C^+^ monocytic population remained largely unchanged with increasing RLT dose (**Figure S3c-e**). F4/80 expression was reduced across all three populations with increasing RLT dose (**Figure S3g-i**). RLT did not significantly alter T cell infiltration across doses (**Figure S3j-k**). However, a trend toward reduced expression of checkpoint markers, including PD-1, TIM-3, and LAG-3, was observed following RLT (**Figure S3l-r**). Collectively, these findings demonstrate that Lu-177 RLT reshapes the prostate TME, particularly within the myeloid compartment.

### Lu-177 RLT combined with Cy promotes proinflammatory myeloid remodeling and enhances endogenous T cells

We previously demonstrated that cyclophosphamide (Cy) pre-conditioning modulates the TME and enhances both endogenous and adoptively transferred CAR T cells (*1*). Specifically, Cy-mediated lymphodepletion promoted pro-inflammatory immune remodeling and improved CAR T cell expansion and tumor infiltration. Given these established immunomodulatory effects of Cy, we sought to determine whether RLT could further reshape the TME beyond the contribution of Cy alone. Here, Cy was administered as previously described, followed by Lu-177 RLT at 37, 74, or 120 MBq (**Figure 3a**). Following Cy pre-conditioning, RLT induced a significant reduction in tumor volume by 6 days post-RLT (P < 0.0166) (**Figure 3b**). In Cy + RLT-treated mice, CD3^+^ tumor-infiltrating lymphocytes (TILs) were increased compared to Cy alone within the CD45^+^ immune compartment (**Figure 3c-d**). Within these T cells, the increase was predominantly driven by CD4^+^ T cells in a dose-dependent manner, whereas the frequency of CD8^+^ T cells remained relatively unchanged (**Figure 3e-f**). We further observed a reduction in the expression of checkpoint markers TIM-3 and LAG-3 but not PD-1, with the most pronounced effect at the highest RLT dose (**Figure 3g-i**). Consistently, the proportion of PD-1^+^ TIM-3^+^ LAG-3^+^ triple-positive T cells, indicative of a more exhausted phenotype, was reduced in RLT-treated mice (**Figure 3j**). Moreover, tumor-infiltrating T cells shifted from a naïve (Tn) phenotype toward an effector-memory/effector-like (Tem/Teff) state, suggesting enhanced functional activation by RLT (**Figure 3k-m**). In parallel, the myeloid compartment exhibited a shift toward granulocytic populations following RLT treatment, consistent with RLT alone (**Figure 3n**).

**Figure 3.**
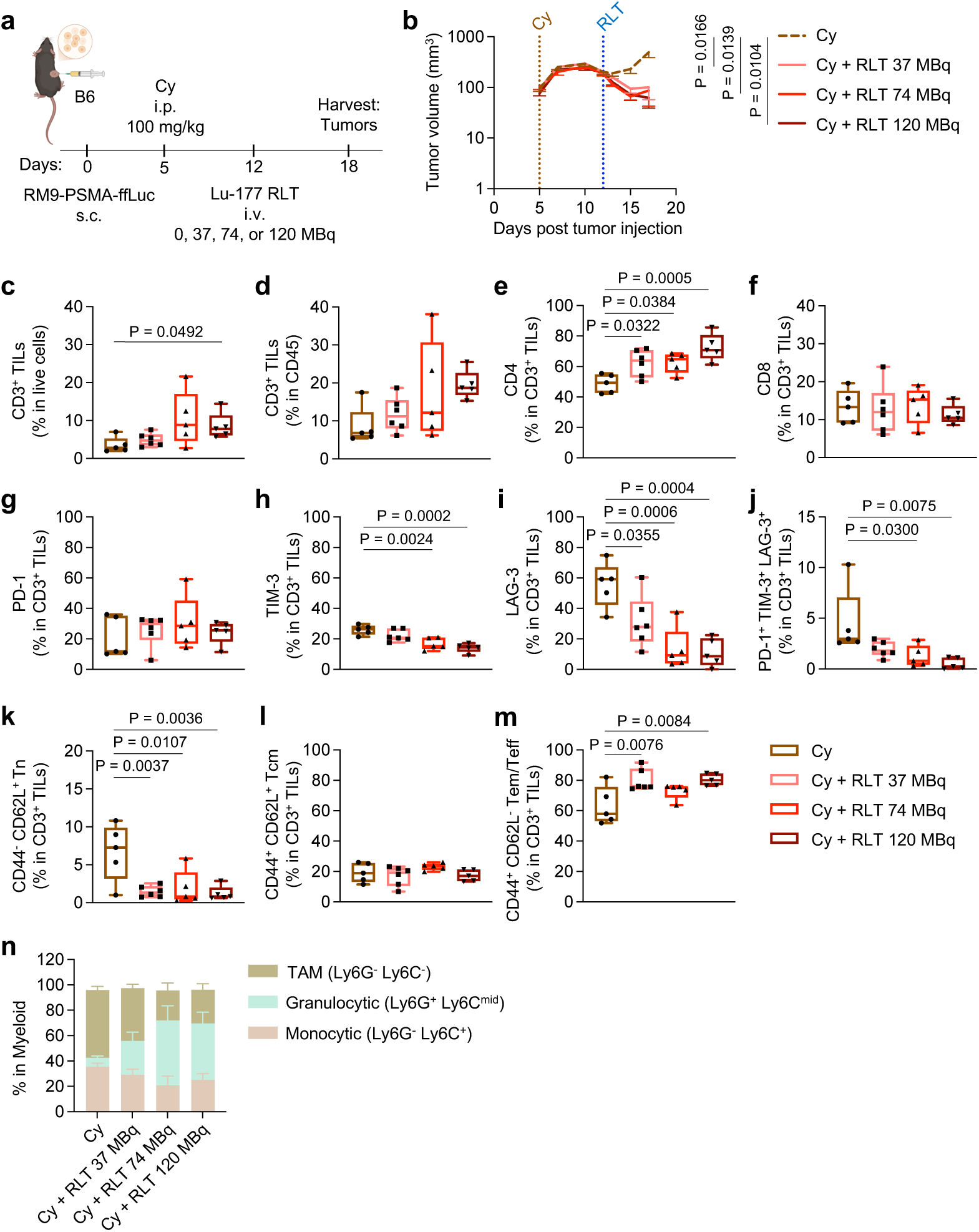
Lu-177 RLT combined with Cy remodels the endogenous T cell phenotypes and myeloid composition in mouse syngeneic prostate cancer models. (**a**) Schematic of tumor injection and treatment schedule in C57BL/6J mice bearing subcutaneous murine prostate cancer. s.c., subcutaneous, i.p., intraperitoneal. On day 5 post-tumor injection, mice received Cy (100 mg/kg, i.p.), followed by Lu-177 RLT (37, 74, or 120 MBq, i.v.) on day 12. Six days later (day 18), mice (n = 5-6 per group) were euthanized, and tumors were harvested for flow cytometric analysis. (**b**) Tumor volume of mice treated with Cy alone, or Cy + Lu-177 RLT at 37, 74, or 120 MBq (n = 5 per group, except Cy + RLT 37 MBq, n = 6). Tumor volume was measured with calipers twice weekly until day 18 (study endpoint). Statistical analysis was assessed by two-way ANOVA with Geisser-Greenhouse correction, followed by Dunnett’s multiple comparison test. (**c-n**) Flow cytometric analysis of tumor single-cell suspensions collected at 6 days post-RLT treatment (n = 5-6 per group). Percentage of CD3^+^ in total live cells (**c**) and CD45^+^ immune cells (**d**); percentage of CD4^+^ (**e**) and CD8^+^ T cells (**f)**; percentage of checkpoint markers PD-1 (**g**), TIM-3 (**h**), LAG-3 (**i**), and PD-1^+^ TIM-3^+^ LAG-3^+^ triple-positive T cells (**j**); percentage of CD44^-^CD62L^+^ naïve T cells (Tn, **k**), CD44^+^ CD62L^+^ central memory T cells (Tcm, **l**), and CD44^+^ CD62L^-^effector memory and effector T cells (Tem/Teff, **m**); and percentage of Ly6G^-^ Ly6C^-^ tumor associated macrophages (TAM), Ly6G^+^ Ly6C^mid^ granulocytic cells, and Ly6G^-^ Ly6C^+^ monocytic cells (gated on CD11b^+^, **n**). Box plots represent median with interquartile range and whiskers indicating minimum and maximum; bar graphs represent mean ± SEM. Statistical significance was determined by one-way ANOVA with Bonferroni correction.

To further investigate the immunomodulatory effects of RLT alone or in combination with Cy, we performed single-cell RNA-sequencing (scRNA-seq) in tumors from untreated (UT), Lu-177 RLT, Cy, or Cy + RLT combination-treated mice 6 days post-RLT treatment (**Figure 4a**). Although 120 MBq demonstrated more pronounced immunomodulatory effects, 74 MBq was selected to preserve renal function over the duration of long-term experiments, as radiation-induced nephrotoxicity in mice manifests in a dose-dependent manner at late timepoints (*19, 20*). Cy was administered on day 9 (3 days post-RLT), aligned with the therapeutic sequence (RLT followed by Cy and PSCA CAR T cells) to both recapitulate the treatment regimen and enable a 4-day interval prior to CAR T cell infusion in subsequent studies. Uniform Manifold Approximation and Projection (UMAP) coupled with ScType-based annotation, identified 10 immune cell populations (**Figure 4b**). Notably, flow cytometry and scRNA-seq datasets provide complementary results in the myeloid cell compartment: the flow-defined Ly6G^−^Ly6C^−^ TAM-like population broadly corresponds to transcriptionally resolved macrophage clusters in scRNA-seq, including M1-like, M2-like, and macrophage-other subsets, whereas the Ly6G^+^Ly6C^mid^ granulocytic and Ly6G^−^Ly6C^+^ monocytic gates correspond predominantly to neutrophil and monocyte populations, respectively. Prior studies have shown that radiation promotes proinflammatory reprogramming of TAMs; therefore, we focused on M1-like macrophages, which exhibited the most pronounced and consistent transcriptional changes in our studies. Within M1-like macrophages, differential gene expression analysis revealed that RLT treatment alone was associated with enrichment of pathways related to immune cell chemotaxis, involving both myeloid and lymphoid compartments, as well as enhanced T cell proliferation and differentiation, compared to UT controls (**Figure 4c**). Upon addition of Cy, M1-like macrophages showed further enrichment in pathways related to antigen processing and presentation (**Figure 4d**). Consistently, heatmap analysis of the significantly differentiated-expressed genes (adjusted P < 0.05) in the combination-treated group highlighted upregulation of genes associated with innate immune activation (*Tlr2*, *Clec5a*, *Clec4d*, *Cd14*, and *Lyz2*), immune cell recruitment (*Ccl4*, *Cxcl16*, *Ccr2*, and *Cxcr4*), and antigen presentation (*H2-Aa*, *H2-Ab2*, *H2-Eb1*, *H2-DMa*, *H2-DMb1*, *Cd74*, and *H2-D1*) (**Figure 4e**). Across MHC class II and I-related genes, we observed that RLT alone modestly upregulates MHC class I (*H2-K1* and *H2-D1*, and non-classical H2-Q4, *H2-T22,* and *H2-T23*). In contrast, the combination resulted in broader upregulation of both MHC class II and I genes (**Figure 4f**).

**Figure 4.**
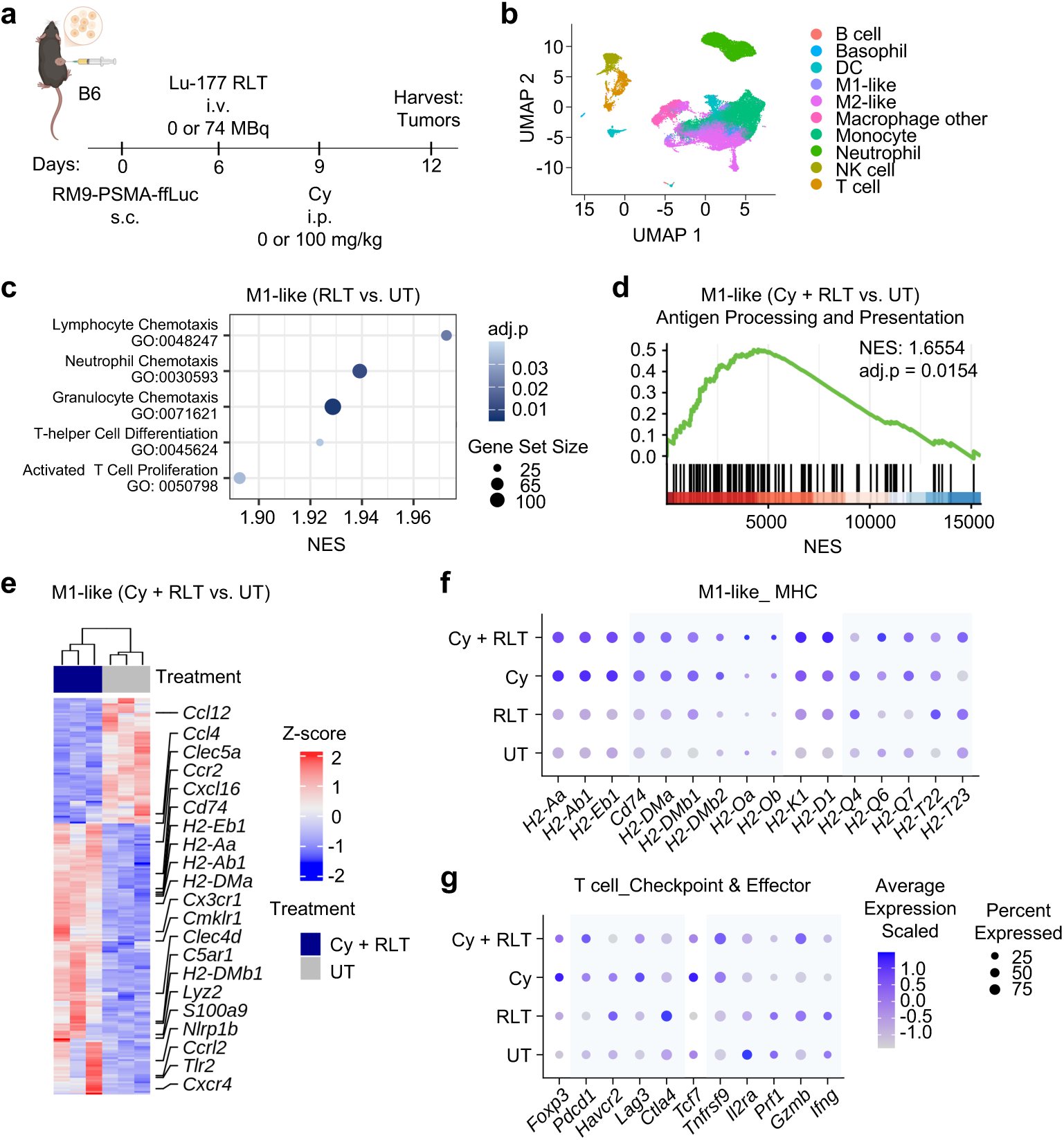
Lu-177 RLT alone or in combination with Cy remodels the prostate TME revealed by scRNA-seq analysis. (**a**) Schematic of tumor injection and treatment schedule in C57BL/6J mice bearing subcutaneous prostate tumors. On day 6 post-tumor injection, mice received Lu-177 RLT (74 MBq, i.v.), followed by Cy (100 mg/kg, i.p.) on day 9. Tumors were harvested 6 days post-RLT (day 12) for scRNA-seq analysis (n = 3 per group). (**b**) UMAP visualization of immune cell populations annotated by ScType, including B cells, basophils, dendritic cells (DCs), macrophages, monocytes, neutrophils, natural killer (NK) cells, and T cells. Cell identities were assigned based on clustering and canonical marker expression. Macrophages were further subdivided into M1-like, M2-like, and macrophage-other subsets. (**c**) Representative top enriched immune-related pathways in M1-like macrophages comparing Lu-177 RLT versus untreated (UT) groups, identified by Gene Set Enrichment Analysis (GSEA) and ranked by normalized enrichment score (NES). (**d**) Representative GSEA enrichment plot for pathway enrichment in M1-like macrophages from Cy + RLT versus UT. (**e**) Heatmap of differentially expressed genes in M1-like macrophages comparing Cy + RLT versus UT (adjusted p < 0.05, |log₂ fold change| > 0.58). Genes shown are macrophage-associated transcripts. (**f**) Balloon plot of MHC class I and II genes (classical and non-classical) in M1-like macrophages across treatment groups (UT, RLT, Cy, and Cy + RLT). Dot size represents the proportion of expressing cells, and color intensity reflects relative expression. (**g**) Balloon plot of checkpoint and effector gene expression in endogenous T cells across treatment groups.

In addition to M1-like macrophages, other immune cell populations also exhibited transcriptional changes following treatment. NK cells from the RLT-treated group showed enrichment in pathways related to antigen presentation via MHC class II (**Figure S4a**), while neutrophils demonstrated negative enrichment for angiogenesis-related pathways, suggesting reduced pro-angiogenic transcriptional activity in this cell population (**Figure S4b**). With the addition of Cy, not only M1-like macrophages but also M2-like and other macrophages (reflecting the diversity of macrophage states beyond the M1/M2 polarization) displayed enrichment in pathways associated with chemotaxis, cell migration, or antigen presentation pathway enrichment (**Figure S4c-f**). NK cells in the Cy + RLT group similarly showed negative enrichment for angiogenesis pathways (**Figure S4g**). Consistent with observations in M1-like macrophages, broader upregulation of both MHC class I and II genes was observed across additional myeloid populations, including M2-like macrophages, macrophage-other subsets, and monocytes (**Figure S5a-c**). In dendritic cells (DCs), RLT alone preferentially upregulated MHC class II genes, whereas the combination of Cy + RLT resulted in greater upregulation of MHC class I genes (**Figure S5d**).

These changes across multiple immune populations within the TME were associated with shifts in endogenous T cell phenotypes. Within the endogenous tumor-infiltrating T cells, we observed upregulation in *Pdcd1*, but not other checkpoint markers, *Havcr2* (encoding TIM-3), *Lag3*, or *Ctla4* (**Figure 4g**). Notably, *Tnfrsf9* (encoding 4-1BB) was also increased at the population level, suggesting that the elevated *Pdcd1* is more consistent with T cell activation than exhaustion. We observed modest upregulation in *Gzmb*, but not the other key effector genes, *Pfr1* or *Ifng*. Collectively, these results show that RLT combined with Cy induced immune activation, thereby providing a rationale for incorporating CAR T cell therapy to further amplify anti-tumor immunity and unlock the potential for durable tumor control.

### Lu-177 RLT and CAR T cell combination treatment enhances antitumor efficacy in mouse syngeneic prostate cancer models

Building on the observed immunomodulatory effects of RLT in immunocompetent settings, we next evaluated the therapeutic efficacy and safety of combining RLT, Cy lymphodepletion, and PSCA-CAR T cells in the syngeneic prostate cancer model. To this end, we employed an antigen-heterogeneous tumor model composed of a mixture of single PSMA-expressing and single PSCA-expressing RM9 tumor cells (**Figure S6a-b**). RM9-PSMA-ffLuc and RM9-PSCA-ffLuc were combined on the day of tumor injection to form a mixed cell suspension, and immediately engrafted subcutaneously in hPSCA knock-in (hPSCA-KI) C57BL/6 mice. The mixture was designed to approximate balanced antigen representation *in vivo* at the time of RLT and CAR T cell treatment. These transgenic mice are tolerant to hPSCA and express the antigen in physiologically relevant tissues, including the bladder and stomach (*21*), enabling assessment of both anti-tumor efficacy and on-target, off-tumor toxicity. Tumor-bearing mice were treated sequentially with Lu-177 RLT (74 MBq), followed 3 days later with Cy (100 mg/kg), and then 1 day later with PSCA-CAR T cells (1 x 10^6^ cells) (**Figure 5a**). TAG72-CAR T cells were used as a non-targeting control. As a positive control, mice received Cy followed by co-administration of both PSCA-CAR T cells and PSMA-CAR T cells (1 x 10^6^ cells each) to demonstrate that antigen-heterogeneous tumors can be effectively treated with dual-targeting CAR T cells. hPSMA and hPSCA antigen expressions were retained on treatment days (day 5 and day 9, corresponding to RLT and CAR T cell administration, respectively) (**Figure S6c**). The individual tumor growth curve showed that combinations of control TAG72-CAR T cells (RLT + TAG72-CAR or Cy + TAG72-CAR) produced only transient anti-tumor efficacy, with rapid tumor progression (**Figure 5b**). The combination of Cy and PSCA-CAR T cells improved tumor control, demonstrating antigen-specific activity. However, despite the initial response, Cy + PSCA-CAR T cell treatment showed tumor regrowth in 8 of 9 mice within 2 weeks, consistent with the absence of PSMA-targeting in this regimen. In contrast, the triple combination (RLT + Cy + PSCA-CAR) achieved a rapid anti-tumor response with a higher curative response rate (5 in 9 versus 1 in 9 with Cy + PSCA-CAR), indicating an added benefit from PSMA targeting with RLT (P = 0.0414) (**Figure 5c**). Moreover, this regimen outperformed the corresponding non-targeting control (RLT + Cy + TAG72-CAR), supporting the requirement for dual-antigen-specific targeting in achieving durable responses. Notably, the therapeutic efficacy of the triple combination was comparable to that of the dual-CAR T cell control. Body weight changes were unremarkable for all treatment groups (**Figure 5d**).

**Figure 5.**
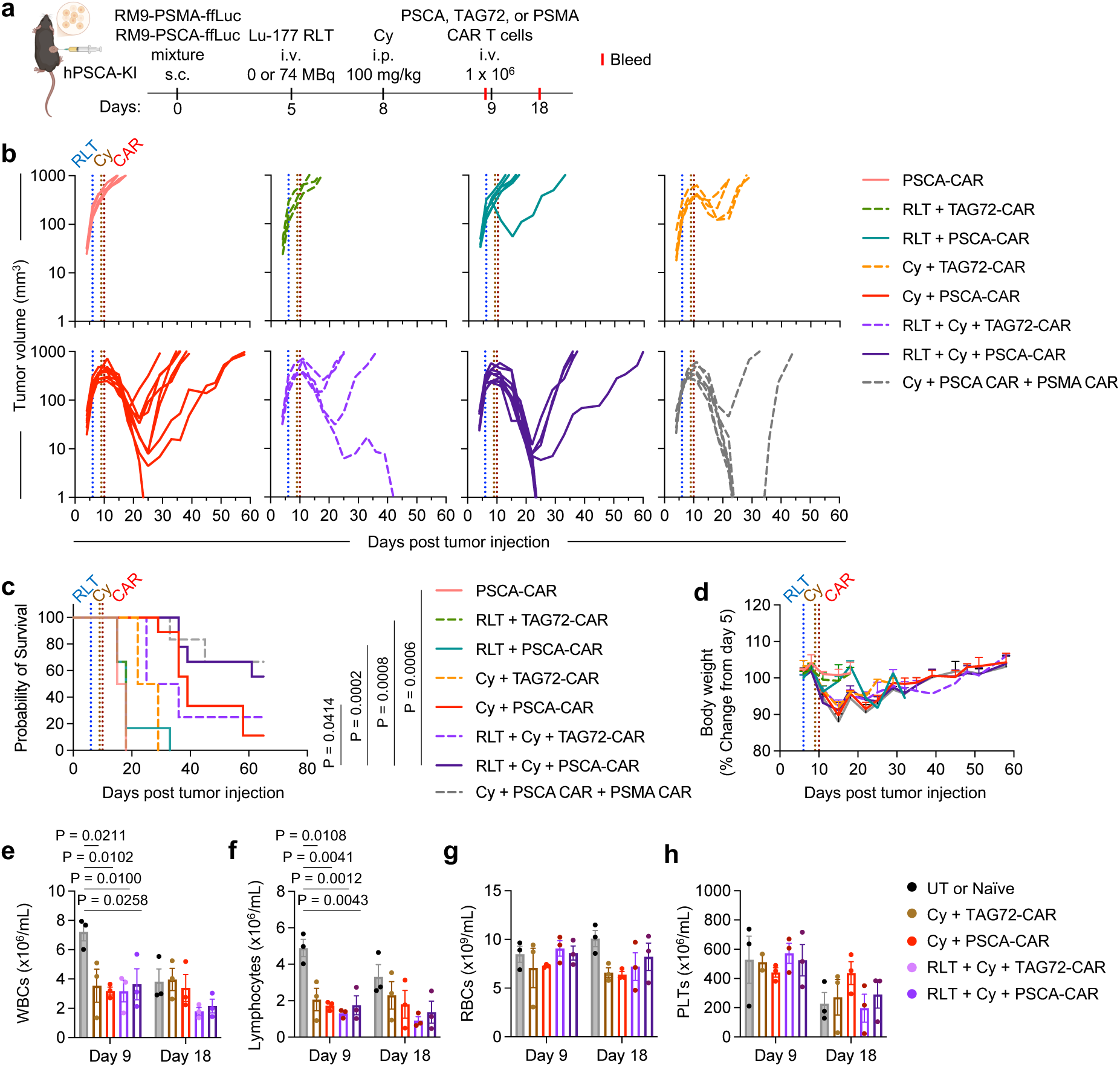
Lu-177 RLT, Cy, and PSCA-CAR T cell combination enhances therapeutic efficacy and survival in syngeneic prostate cancer models. (**a**) Schematic of tumor injection and treatment schedule in hPSCA-KI C57BL/6J mice bearing antigen-heterogeneous subcutaneous prostate tumors. s.c. subcutaneous; i.v., intravenous. RM9-PSMA-ffLuc and RM9-PSCA-ffLuc were mixed prior to tumor injection, and 1.5 × 10^6^ total cells were engrafted into the left flank. On day 5 post-tumor injection, mice received Lu-177 RLT (74 MBq, i.v.), followed by Cy (100 mg/kg, i.p.) on day 8, and CAR T cells (PSCA-CAR, non-targeting TAG72-CAR control, or PSMA-CAR positive control, 1 × 10^6^, i.v.). The positive control group received 2 × 10⁶ total CAR T cells (1 × 10⁶ of each CAR). Blood samples were collected on day 9 (prior to CAR T cell infusion) and day 18 (tumor regression phase) for CBC analysis (n = 3 per group). (**b**) Individual tumor growth curves. Tumor volume was measured by caliper twice weekly until the survival endpoint (n = 9 for RLT + Cy + PSCA-CAR/TAG72-CAR; n = 6 for RLT + PSCA-CAR and Cy + PSMA-CAR + PSCA-CAR; n = 4 for PSCA-CAR, RLT + PSCA-CAR, and Cy + TAG72-CAR; and n = 3 for RLT + TAG72-CAR). (**c**) Kaplan-Meier survival curves for each treatment group. P-value is calculated using multiple comparisons with Holm-Šídák’s correction. (**d**) Body weight changes over time, normalized to baseline (day 5, prior to RLT). Data are presented as mean ± SEM. (**e-h**) Hematologic analysis by CBC, including white blood cells (WBCs, **e**), lymphocytes (**f**), red blood cells (RBCs, **g**), and platelets (PLTs, **h**) from each group (n = 3 per group). Controls include tumor-bearing untreated (UT) mice at day 9, and tumor-naïve mice at day 18. Data are presented as mean ± SEM. Statistical significance was assessed using two-way ANOVA, followed by Tukey’s multiple comparison test.

To assess safety, blood samples were collected on day 9 (prior to CAR T cell infusion) and day 18 (during tumor regression). A consistent reduction in WBCs and lymphocytes was observed in all treatment groups at day 9 compared to control, consistent with the lymphodepletion effect from Cy. (**Figure 5e-f**). By day 18, the counts were comparable to tumor-naïve controls. RBCs, PLTs, and hemoglobin (HGBs) levels were at comparable levels across all groups at both timepoints showing absence of anemia or thrombocytopenia related to treatment (**Figure 5h, S7a**). Serum chemistry analysis showed no significant changes in markers of renal or hepatic function, including blood urea nitrogen (BUN), creatinine, alanine aminotransferase (ALT), and aspartate aminotransferase (AST), except elevation of creatinine in the Cy + PSCA-CAR group (**Figure S7b-e**). Histological evaluation of major organs, including lung, liver, stomach, intestine, kidney, and bladder, revealed no overt tissue damage by H&E staining across treatment groups (**Figure S7f**). Collectively, these data support the combination of Lu-177 RLT, Cy, and PSCA-CAR T cells in improving anti-tumor efficacy and survival in the absence of treatment-related toxicity.

We next assessed whether mice cured with RLT + Cy + PSCA-CAR T cell therapy developed immunological memory. On day 123 post tumor injection (∼2 months after confirmed tumor eradication), mice were rechallenged subcutaneously with 0.5 × 10^6^ RM9-WT cells (lacking PSMA or PSCA expression). Compared to age-matched, treatment-naïve controls, previously cured mice exhibited delayed tumor growth following rechallenge (**Figure S8a**). Notably, tumor inhibition was observed despite the absence of target antigen expression, suggesting the prior therapy may have induced broader anti-tumor immune responses. Among non-cured mice treated with RLT + Cy + PSCA-CAR, early (≤40 days) and late (∼60 days) relapse patterns were observed (**Figure 5b**). To investigate potential resistance mechanisms, recurrent tumors were analyzed by IHC for PSMA and PSCA expression. Early relapsing tumors retained PSMA but lacked PSCA expression, suggesting effective elimination of PSCA^+^ tumor cells by CAR T cells, with residual disease likely driven by PSMA-expressing tumor cells (**Figure S8b**). Late relapses from the Cy + PSCA-CAR group similarly retained PSMA expression, whereas tumors from the RLT + Cy + PSCA-CAR group lacked both PSMA and PSCA, consistent with antigen loss or outgrowth of antigen-negative clones as a potential resistance mechanism.

Given that PSMA and PSCA expression are not strictly mutually exclusive, we further evaluated the combination regimen in an antigen-homogeneous syngeneic model (**Figure S9a**). RM9-PSMA-PSCA-ffLuc cells were implanted in hPSCA-KI mice, followed by treatment as described above, with reduced Cy (50 mg/kg) and CAR T cell dose (0.5 × 10^6^ cells). Consistent with prior findings, RLT + Cy + PSCA-CAR T cell treatment induced a more rapid tumor response compared to Cy + PSCA-CAR alone (mean peak tumor burden at day 11 versus day 15, respectively), although the survival benefit did not reach statistical significance (**Figure S9b-c**). All treatments were well tolerated, with no significant changes in body weight observed (**Figure S9d**). Collectively, these data suggest that RLT priming enhances CAR T cell efficacy and overcomes antigen heterogeneity, prompting us to further investigate the underlying cellular and molecular mechanisms.

### Lu-177 RLT enhances PSCA-CAR T cell activation and effector programs while promoting proinflammatory myeloid remodeling

To investigate mechanisms underlying enhanced tumor control and survival observed with the combination strategy, we performed scRNA-seq on tumors treated with Cy + PSCA-CAR or control TAG72-CAR T cells, with or without the combination of RLT. Tumors were harvested on day 18 (13 days post-RLT and 9 days post-CAR T cell infusion) (**Figure 6a**). We were able to detect CAR T cells, either PSCA-CAR or TAG72-CAR, in tumors (**Figure 6b**). Comparing RLT + Cy + PSCA-CAR group with the Cy + PSCA-CAR group, CAR T cells exhibited increased expression of activation markers (*Tnfrsf9* and *Il2ra*) and effector genes most notably *Ifng* (**Figure 6c**). Differential gene expression analysis further revealed upregulation of genes associated with activation (*Tnfsf4*, *Tnfsf13*, *Fos*, and *Ltbr*), stemness (*Tcf7*), TCR signaling (*Hck*, *Pik3cb*), and effector function (*Batf2*, *Cd63*, and *Ifi30*) (**Figure 6d**). Pathway enrichment analysis of CAR T cells suggested that RLT may enhance CAR T cell programs related to cytokine production, activation, and chemotaxis, particularly toward myeloid populations (**Figure 6e**). M1-like macrophages in the RLT + Cy + PSCA-CAR group showed enrichment of pathways associated with immune cell chemotaxis, extending our earlier observations of RLT- and Cy + RLT-mediated myeloid remodeling (**Figure 6f, 3n, & 4c**). Additionally, interferon-related pathways and cytokine signaling associated with CD4^+^ T cell regulation were observed. Of note, there was an enrichment within macrophages and monocyte populations with respect to interferon pathways and cytokine production important for CD4 T cells regulation, suggesting coordinated crosstalk between T cell and myeloid cell compartments in response to the combination therapy (**Figure S10a-b**). To further investigate inter-cellular interactions, we performed cell-cell communication analysis. Comparing signaling information flow from M1-like macrophages to CAR T cells, we observed near-complete dominance of MHC-II, MIF, and ADGRL signaling in the RLT + Cy + PSCA group, indicating enhanced antigen presentation and inflammatory communication toward CAR T cells (**Figure 6g**). In contrast, the Cy + PSCA group showed higher relative contribution from CD200 and CD48 pathways, which are associated with immune regulation and suppressive signaling, in the absence of RLT. Beyond M1-like macrophages, we also observed enrichment of IL16 and TNF ligand-receptor interactions originating from M2-like macrophages and monocytes towards CAR T cells, further supporting the coordinated network of myeloid-CAR T cell communication that may contribute to the enhanced efficacy of the combination therapy (**Figure S10c-d**). Collectively, these findings suggest that RLT enhances CAR T cell activation and coordinates proinflammatory myeloid cell responses.

**Figure 6.**
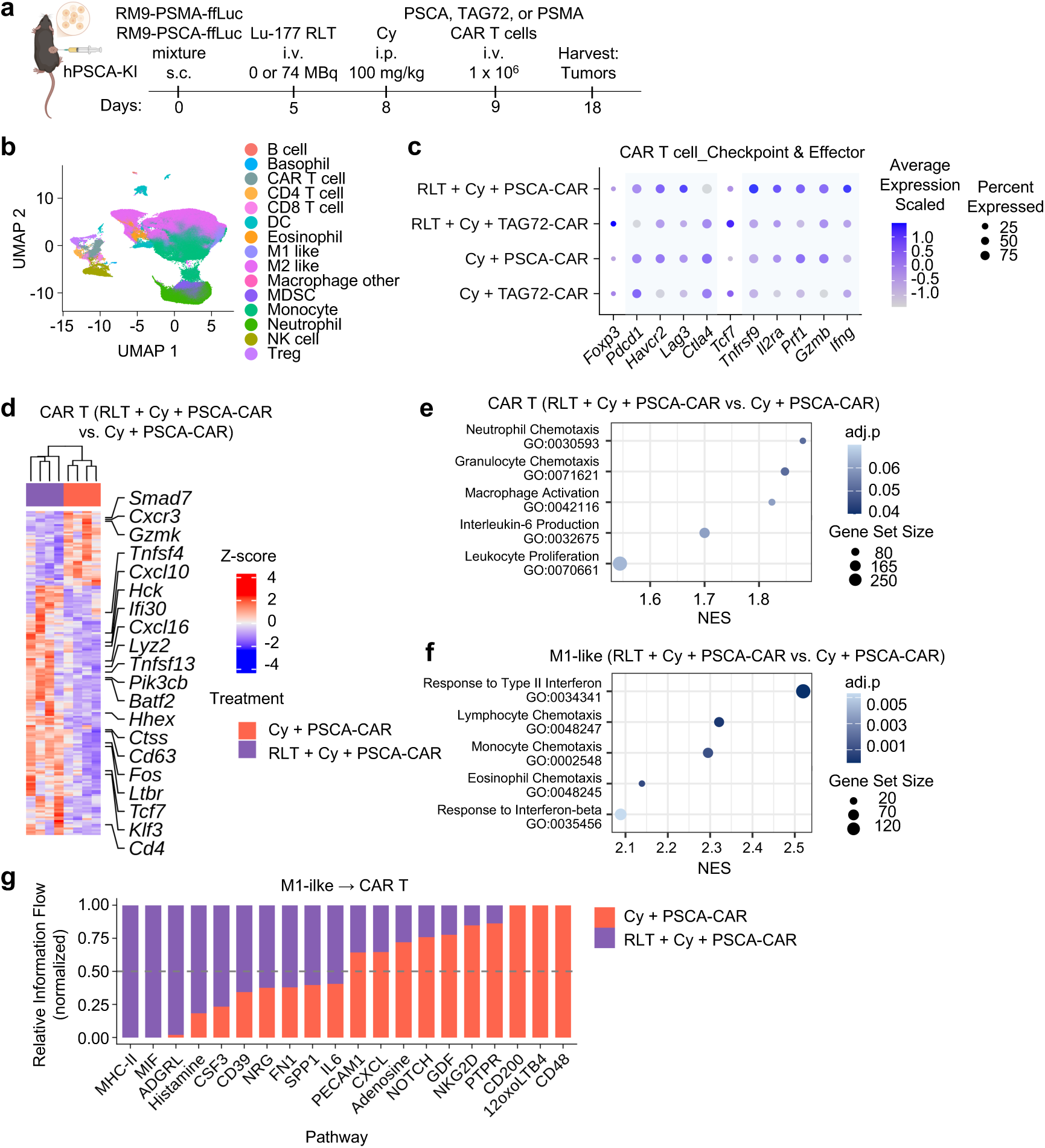
Lu-177 RLT enhances PSCA-CAR T cell activation and effector programs via myeloid cell communication. (**a**) Schematic of tumor injection and treatment schedule in hPSCA-KI C57BL/6J mice bearing antigen-heterogeneous subcutaneous prostate tumors. 1.5 × 10^6^ total tumor cells were injected into the left flank. On day 5 post-tumor injection, mice received Lu-177 RLT (74 MBq, i.v.), followed by Cy (100 mg/kg, i.p.) on day 8, and CAR T cells (PSCA-CAR or TAG72-CAR control, 1 × 10^6^, i.v.). Tumors were harvested 9 days post-CAR T cell treatment (day 18) for scRNA-seq analysis (n = 4 per group). (**b**) UMAP visualization of immune cell populations annotated by ScType. Major immune cell types, including B cells, basophils, T cells, dendritic cells (DCs), eosinophils, macrophages, monocytic-derived suppressor cells (MDSCs), monocytes, neutrophils, and M1-like, M2-like, and macrophage-other subsets. T cells were further subdivided into endogenous regulatory T cells (Tregs), endogenous CD4 (excluding Tregs), endogenous CD8, and adoptively transferred CAR T cells. (**c**) Balloon plot of checkpoint and effector gene expression in CAR T cells across treatment groups (Cy + TAG72-CAR, Cy + PSCA-CAR, RLT + Cy + TAG72-CAR, and RLT + Cy + PSCA-CAR). (**d**) Heatmap of differentially expressed genes in CAR T cells comparing RLT + Cy + PSCA-CAR versus Cy + PSCA-CAR (adjusted p < 0.05, |log₂ fold change| > 0.58). Genes shown are T cell-associated transcripts. (**e-f**) Representative top enriched immune-related pathways in CAR T cells (**e**), and M1-like macrophages (**f**), comparing RLT + Cy + PSCA-CAR versus Cy + PSCA-CAR groups, ranked by normalized enrichment score (NES). (**g**) Cell-cell communication analysis by CellChat, showing signaling interactions from natural killer (NK) cells, were defined based on clustering and canonical marker expression. Macrophages were further subdivided into M1-like macrophages (ligand) to CAR T cells (receptor). Bar plots represent relative information flow, normalized across conditions, with the top 10 pathways shown for each group.

### Combination of Ac-225 RLT with CAR T cells enhances anti-tumor efficacy in prostate cancer

While beta-emitter Lu-177 has gained FDA approval, increasing attention has been directed toward the alpha-emitter Ac-225 as a promising RLT strategy for treating mCRPC (*22*). ^225^Ac-PSMA-617 (Ac-225 RLT) delivers a high radiation dose per decay and is particularly effective against small-volume disease, such as micrometastases and circulating tumor cells (*23*). In the metastatic human xenograft model, NSG mice were injected intracardially with PC3-PIP-PSCA-ffLuc cells, followed by Ac-225 RLT treatment (15 kBq), and PSCA-CAR T cells (0.2 x 10^6^) administered 4 or 6 days after RLT (**Figure 7a**). Ac-225 RLT monotherapy resulted in minimal tumor control, whereas combination with PSCA-CAR T cells produced greater control and a more durable anti-tumor response (**Figure 7b**). Survival analysis demonstrated that the combination significantly improved survival compared to RLT alone (P = 0.0004), with a trend toward improved survival compared to PSCA-CAR T cell monotherapy (P = 0.0952) (**Figure 7c**). We further evaluated this combination in the antigen-heterogeneous syngeneic prostate cancer model. hPSCA-KI C57BL/6 mice bearing a mixture of RM9-PSMA-ffLuc and RM9-PSCA-ffLuc tumors were treated with Ac-225 RLT (18.5 kBq), followed 3 days later with Cy (100 mg/kg), and 1 day later with PSCA-CAR or control TAG72-CAR T cells (1 x 10^6^) (**Figure 7d**). Consistent with prior observations, PSCA-CAR T cells alone induced transient tumor control (**Figure 7e**). The addition of Ac-225 RLT as a priming strategy improved tumor control. We observed a trend toward a higher proportion of responders in the RLT + Cy + PSCA-CAR group compared to the non-targeting CAR T cell control, supporting the importance of antigen-specificity. Survival analysis demonstrated that the combination strategy significantly improved overall survival compared to RLT (P = 0.0015) or PSCA-CAR T cell monotherapy (P = 0.0492). Although the overall body weight change did not exceed the 20% threshold, we observed treatment-associated weight loss in groups receiving Ac-225 RLT in combination with Cy, independent of CAR T cell treatment (**Figure 7g**). Collectively, these findings demonstrate that alpha-emitter RLT enhances anti-tumor activity in combination with CAR T cells, while further optimization of Cy dosing may be warranted to improve tolerability in combination regimens for future clinical development.

**Figure 7.**
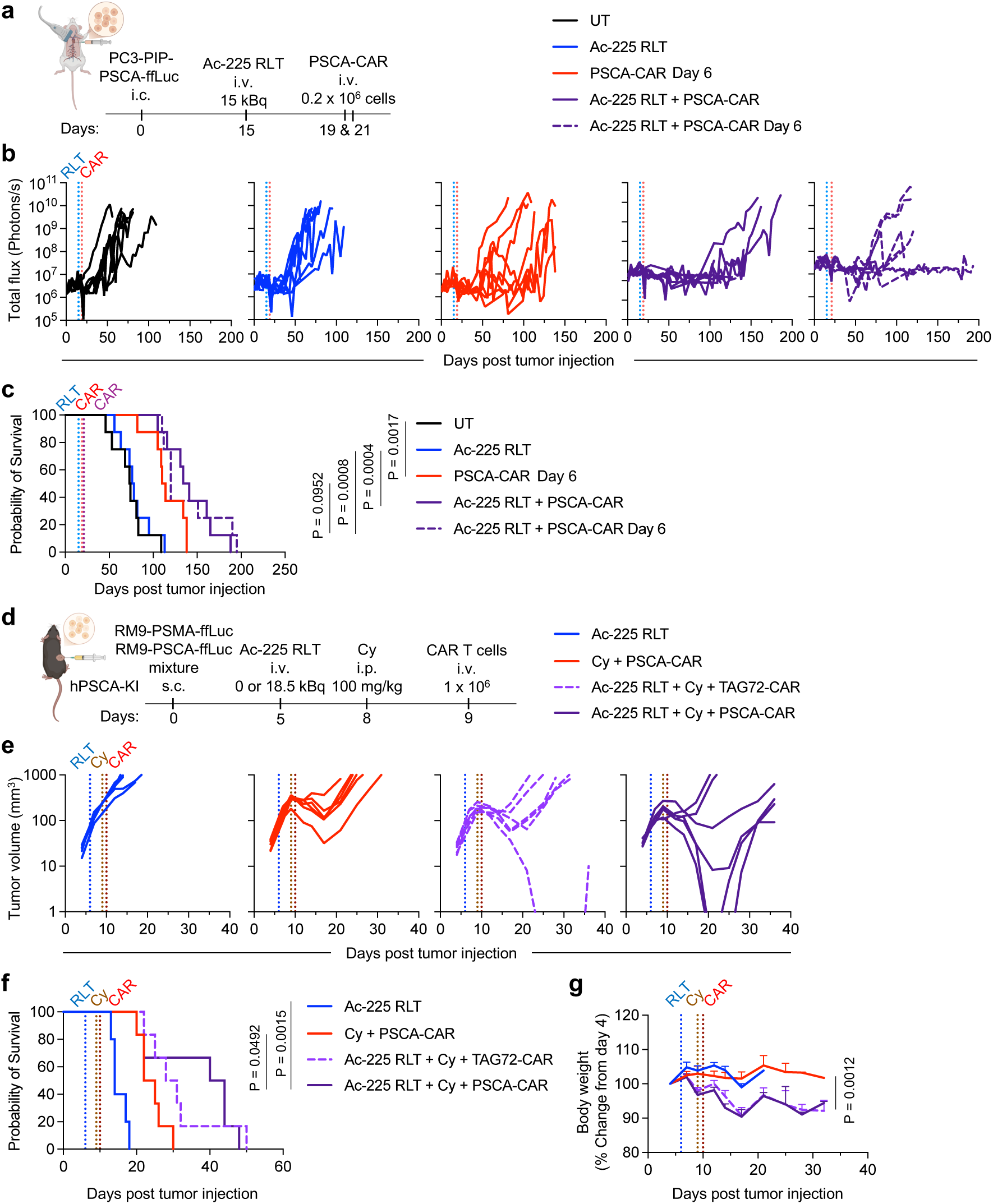
Ac-225 RLT and PSCA-CAR T cell combination enhances anti-tumor response in human xenograft and mouse syngeneic prostate cancer models. (**a**) Schematic of tumor injection and treatment schedule in NSG mice bearing metastatic prostate cancer. On day 15 post-tumor injection, mice received Ac-225 RLT at 15 kBq (i.v.); followed by PSCA-CAR T cells (0.2 × 10^6^, i.v.) 4 or 6 days later (day 19 or 21). PSCA-CAR T cell monotherapy was administered on day 21. (**b**) Longitudinal whole-body BLI of PC3-PIP-PSCA-ffLuc metastatic tumor-bearing mice treated with Ac-225 RLT alone, PSCA-CAR T cells alone, or the combination (n = 8 per group). BLI was performed twice a week until day 193, and quantified using a consistent ROI. (**c**) Kaplan-Meier survival curves. P-value is calculated using multiple comparisons with Holm-Šídák’s correction. (**d**) Schematic of tumor injection and treatment schedule in hPSCA-KI C57BL/6J mice bearing antigen-heterogeneous subcutaneous tumors. 1.5 × 10^6^ total cells were engrafted into the left flank. On day 5 post-tumor inoculation, mice received Ac-225 RLT (18.5 kBq, i.v.), followed by Cy (100 mg/kg, i.p.) on day 8, and CAR T cells (PSCA-CAR or TAG72-CAR, 1 × 10^6^, i.v.). (**e**) Individual tumor volume of each group (n = 6 per group, except n = 5 for Ac-225 RLT alone). (**f**) Kaplan-Meier survival curves for each treatment group (Holm-Šídák’s correction). (**g**) Body weight changes over time in each group, normalized to baseline (day 5, pre-RLT). Data are presented as mean ± SEM. Statistical significance was assessed by unpaired t-test with multiple comparisons correction. P-value shown corresponds to comparisons between the Cy + PSCA-CAR and RLT + Cy + PSCA-CAR groups at day 14 post tumor injection.

## DISCUSSION

In this study, we evaluated the efficacy and safety of combining RLT with CAR T cell therapy across multiple prostate cancer models. This approach was designed to address two major barriers limiting current treatments in prostate cancer: tumor antigen heterogeneity and the immunosuppressive TME. In metastatic human xenograft models, the combination demonstrated anti-tumor activity and tolerability in clinically relevant immunodeficient settings that partially recapitulate aspects of mCRPC. These findings were validated in immunocompetent mouse models, where antigen-heterogeneous tumors were more effectively controlled by the combination of RLT, Cy, and CAR T cells. Mechanistically, Lu-177 RLT, particularly in combination with Cy, reprogrammed the tumor myeloid compartment toward a proinflammatory state characterized by enhanced antigen presentation programs and chemokine-mediated recruitment pathways. These changes were associated with activation of endogenous T cells and enhanced CAR T cell activation, effector gene expression, and cytokine signaling. Together, these findings identify myeloid cell-T cell crosstalk as potential mechanism by which RLT primes the TME to support adoptive cell therapy and sustain anti-tumor responses. Importantly, this strategy integrates clinically relevant components, including PSMA-targeted Lu-177 RLT, an FDA approved therapy for mCRPC, and PSCA-targeted CAR T cells with demonstrated safety and bioactivity in phase 1 clinical trials. These features support the clinical translatability of this combination approach. In addition, we evaluated the alpha-emitter Ac-225 RLT as an alternative radioisotope and observed enhanced anti-tumor activity at lower dose in combination with CAR T cells. As clinical experience expands with Ac-225-based, or other radioisotope-based RLT, these insights may inform the development of next-generation combination strategies with radiopharmaceuticals and CAR T cell therapies.

Herein, we employed both human xenograft and mouse syngeneic preclinical models to capture different aspects of prostate cancer biology. However, these models only partially reflect the complexity of antigen expression observed in human disease. Analyses of patient-derived xenografts and public datasets indicate that PSMA and PSCA expressions are highly heterogeneous, with tumors often exhibiting variable or discordant expression rather than the uniformly high co-expression seen in engineered PC3 models or the non-overlapping expression in the RM9 mixture models. Moreover, PSMA and PSCA are predominantly associated with adenocarcinoma, and their expression is frequently downregulated in neuroendocrine prostate cancer (NEPC, more commonly treatment emergent), which accounts for up to 25-30% of mCRPC (*24–26*). Although we observed co-expression of these antigens in tumors with mixed adenocarcinoma/neuroendocrine features, a substantial fraction of tumor cells may lack both targets, representing a potential limitation of this dual-targeting strategy. We are currently investigating the combination strategy incorporating metastasis-directed radiation therapy, cyclophosphamide, and PSCA-CAR T cells in a phase 1b clinical trial (NCT05805371), but in advancing PSMA-directed RLT to clinical investigation, the expression patterns of these two targets may need significant consideration.

Lessons from combining RLT or external beam radiation therapy (EBRT) with immunotherapy highlight that treatment sequence, dosing, and timing are critical determinants of efficacy. Radiation administered after CAR T cell infusion has been associated with inferior outcomes compared to its use as a priming strategy (*27*). Non-ablative doses of ionizing radiation induce immunogenic cell death, enhance antigen presentation, and promote proinflammatory response (*28–30*). This evidence supports the design in which RLT precedes adoptive cell transfer. Although radiation within the tumor may partially impair CAR T cell viability (*31, 32*), a subablative dosing of radiation allows surviving CAR T cells to expand within a primed, cytokine-rich microenvironment, thereby promoting anti-tumor responses.

The choice of radioisotope also influences immune priming, with alpha-emitters such as Ac-255 potentially inducing more cytotoxic and immunomodulatory effects than beta-emitters (*33*). Regarding the timing, CAR T cells require several days to traffic to tumor sites, typically reaching peak accumulation within 1–2 weeks post-infusion. In this study, we administered RLT 4-days prior to CAR T cell infusion, based on our observations that PSCA-CAR T cells require approximately 3-4 days to reach the tumor site (*21*), coinciding with the partial decay of Lu-177 RLT (physical half-life of approximately 6.65 to 6.73 days). Extending the interval from 4 to 6 days did not significantly alter therapeutic efficacy in human xenograft models, although longer intervals could not be evaluated in the mouse syngeneic models due to rapid tumor growth kinetics. In addition to timing, tumor vasculature may further influence the outcomes, as reduced angiogenesis can limit CAR T cell trafficking despite improving tumor control. Accordingly, functional approaches, such as longitudinal tracking of CAR T cell expansion and localization using bioluminescence imaging or CD8-targeted immuno-PET, may provide a more direct means to optimize treatment scheduling (*34*).

Safety remains a key consideration for clinical translation. In our models, no overt toxicities were observed. Especially in our hPSCA-KI syngeneic model that may reflect toxicities associated with targeting PSCA and immune-related events, the combination was well-tolerated. However, potential risks may include on-target, off-tumor effects (e.g., salivary gland and renal toxicity), Cy-and PSCA-CAR T cell-associated bladder toxicity, and RLT-induced cytopenias. These effects may overlap, increasing the risk of immunosuppression, organ damage, and/or cytokine release syndrome. Careful optimization of dose, sequence, and timing will be essential to balance safety and efficacy.

In summary, our study demonstrates that combining RLT with CAR T cell therapy can overcome key barriers to the effective treatment of prostate cancer, including antigen heterogeneity and the immunosuppressive TME. By leveraging the immunomodulatory effects of RLT in priming the immune system in tumors, this approach enhances CAR T cell activity and promotes more durable anti-tumor responses across preclinical human xenograft and mouse syngeneic models. While further optimization may be required, these findings provide a strong foundation for clinical translation of RLT and CAR T cell combination strategies for the treatment of mCRPC and other solid tumor malignancies.

## MATERIALS AND METHODS

### Study design

This study was designed to evaluate the therapeutic efficacy and safety of combining RLT and CAR T cells in preclinical models of prostate cancer. Experiments were performed using both xenograft and immunocompetent syngeneic mouse models with homogenous or heterogeneous antigen expression. All in vivo studies were performed using 5-8 week old NSG or hPSCA-KI C57BL/6 mice, using 3-10 mice included within each group for all therapy and survival studies, and at least three mice per group for *ex vivo* assessment to ensure statistical power. In xenograft studies, NSG mice bearing metastatic tumors were treated with Lu-177 or Ac-225 RLT, followed by PSCA-CAR T cell infusion. In syngeneic studies, hPSCA-KI C57BL/6 mice bearing subcutaneous RM9-derived tumors received Lu-177 or Ac-225 RLT, followed by Cy lymphodepletion and CAR T cells. Before RLT and CAR T cell treatment, mice were randomized based on bioluminescence imaging or tumor volume to ensure an evenly distributed average tumor burden across each group. Treatment groups were compared to appropriate controls, including untreated mice, CAR T cells alone, or RLT/Cy with non-targeting CAR T cells in combination. Animal survival was based on the endpoints (tumor volume >1000mm^3^ or ulceration, or signs of distress including weight loss and impaired mobility). Mechanistic investigation included immune profiling by flow cytometry and scRNA-seq, as well as safety assessments including body weight, hematologic parameters, serum chemistry, and histopathology. Investigators were not blinded to group allocation.

### Cell lines

The PSMA-transfected PC3-PIP cell line was a kind gift from Martin G. Pomper (Johns Hopkins University). Cells were maintained in RPMI-1640 media (Gibco) supplemented with 10% fetal bovine serum (FBS, SeraPrime) and 1% penicillin-streptomycin-glutamine (Gibco). The Ras/Myc transformed murine prostate cancer line RM-9 was a kind gift from Timothy C. Thompson (MD Anderson Cancer Centre). The cell line was cultured in Dulbecco’s Modified Eagle’s Medium (DMEM, Life Technologies) supplemented with 10% fetal bovine serum (FBS, Hyclone), 25 mM HEPES (Irvine Scientific), and 2 mM L-Glutamine (Corning). Cells were detached using 0.25% trypsin (Gibco) for up to 5 min at 37°C for passages and prepared for intracardiac injection in 1X phosphate-buffered saline (PBS, Gibco), or subcutaneous injection in HBSS (Gibco).

### DNA constructs and tumor cell transduction

Tumor cells were engineered to express firefly luciferase (ffLuc), human PSMA, and human PSCA as previously described (*35, 36*). Briefly, PC3-PIP cells were sequentially transduced with epHIV7 lentivirus encoding hPSCA and ffLuc genes, each under the EF1α promoter. RM9 derivatives were generated as follows: RM9-PSCA-ffLuc transduced with hPSCA and ffLuc lentivirus (*21*); RM9-PSMA-ffLuc transduced with hPSMA retrovirus (pMVs backbone, MSCV promoter) and ffLuc lentivirus; RM9-PSMA-PSCA-ffLuc generated by transducing RM9-PSCA-ffLuc with hPSMA retrovirus. All RM9 derivatives were single-cell cloned to ensure homogenous antigen expression.

The PSCA-CAR and non-targeting TAG72-CAR constructs were previously described (*37*). Briefly, the PSCA-CAR consists of murine anti-PSCA scFv (1G8) with murine CD8 hinge, murine CD8 transmembrane domain, murine 4-1BB costimulatory domain, and murine CD3ζ cytoplasmic domain. The TAG72-CAR consists of anti-TAG72 scFv (CC49) and a similar CAR backbone to PSCA-CAR. The CAR construct, followed by the T2A ribosomal skip sequence and truncated murine CD19 tag (CD19t), was cloned into the pMYs retrovirus backbone under the control of a hybrid MMLV/MSCV promoter (Cell BioLabs).

### Viral production

Lentivirus and retrovirus were produced as previously described (*35, 36*). Lentivirus was generated using 293T cells. Briefly, cells were seeded in T-225 tissue culture flasks 1 day before transfection, followed by calcium phosphate transfection using packaging plasmids and the desired CAR-encoding lentiviral backbone plasmid. Supernatants were collected at 72 hours, filtered, and centrifuged to remove cell debris, and incubated with 2 mM magnesium and 25 U/ml Benzonase endonuclease (EMD Millipore) to remove contaminating nucleic acids. Virus-containing supernatants were combined and concentrated via high-speed centrifugation (6,080 × g) overnight at 4 °C. Lentiviral pellets were then resuspended in PBS-lactose solution (4 g lactose per 100 ml PBS), aliquoted, and stored at −80 °C. Lentiviral titers were determined by transducing HT1080 cells.

Retrovirus was generated using the ecotropic retroviral packaging cell line, PLAT-E (Cell Biolabs). Briefly, cells were transfected using FuGENE HD transfection reagent (Promega) with the addition of murine PSCA-CAR or TAG72-CAR encoding retrovirus backbone plasmid DNA. Viral supernatants were collected after 24, 36, and 48 h, pooled, aliquoted, and stored at −80 °C for future T cell transductions.

### Human and mouse CAR T cell generation, cryopreservation, and thawing

Human T cell isolation, PSCA-CAR transduction, expansion, and cryopreservation were previously described (*35*). Murine CAR T cells were generated as previously described (*21*). Briefly, splenocytes were obtained from male hPSCA-KI mice (6-8 weeks old) by mechanical dissociation of spleens in 1X PBS (FUJIFILM Irvine Scientific). Mouse T cells were isolated using EasySep^TM^ mouse T cell negative selection kits according to the manufacturer’s protocol (StemCell Technologies). Mouse T cells were activated with CD3/CD28 Dynabeads (Invitrogen) at a 1:1 ratio, and maintained in RPMI-1640 medium supplemented with 10% fetal bovine serum (FBS, Hyclone), recombinant human/murine IL-2 (50 U/mL; Novartis Oncology), murine IL-7 (10 µg/mL; PeproTech), and 50 µM 2-mercaptoethanol (Gibco). Mouse T cells were cultured for 24 hours, followed by retroviral transduction of CAR constructs using RetroNectin (10 µg/mL, Takara). On day 5, CAR expression was assessed by flow cytometry via CD19t surface expression. CAR T cells were then subjected to Dynabead removal and cryopreserved in FBS containing 10% dimethyl sulfoxide (DMSO, Sigma-Aldrich). On the day of treatment, CAR T cells were thawed in 1X PBS, washed once, and resuspended in 1X PBS at the desired concentration prepared for treatment.

### Radiosynthesis of Lu-177 RLT and Ac-225 RLT

[^177^Lu]Lu-PSMA-617 was used for human xenograft studies. For syngeneic studies, the radiolabeling was performed according to previously published procedures (*38*). Briefly, PSMA-617 precursor (Vipivotide tetraxetan, MedChemExpress) was stored in aliquots in 0.1% aqueous trifluoroacetic acid until use. The radiolabeling was performed by City of Hope Radiopharmacy. The non-carrier-added [^177^Lu]LuCl_3_ (Monrol) was incubated with the precursor in 0.4M acetate buffer (pH 4.5) at 95 °C for 15 min. The specific activity of Lu-177 RLT was ∼60 MBq/nmol. The radiochemical purity was confirmed by instant thin-layer chromatography (iTLC) and high-performance liquid chromatography (HPLC) at ≥ 99%.

For Ac-225 RLT, [^225^Ac]Ac(NO_3_)_3_ (Oak Ridge National Laboratory) was dissolved in 0.1M HCl before use. The radiolabeling was performed by incubating [^225^Ac]Ac(NO_3_)_3_ with the precursor in 0.25M ammonium acetate buffer containing dihydroxiybenzoic acid. The mixture was heated at 95 °C for 30 min, followed by formulation in saline upon cooling. The specific activity is ∼1 MBq/nmol. The radiochemical purity was confirmed by iTLC at ≥ 95%.

### In vitro cell binding and ex vivo biodistribution

For *in vitro* binding assay, cells (RM9-WT, RM9-PSMA-ffLu, RM9-PSCA-ffLuc, and RM9-PSMA-PSCA-ffLuc) were seeded 2-3 days prior to the assay. Cells were detached using 0.25% trypsin, washed once with 1X PBS, and resuspended in HBSS at the desired concentration (1 x 10^7^ /mL for RM9-PSMA-PSCA-ffLuc, or 1.4 x 10^7^ /mL for the other cell lines). Cells were prepared in serial 1:2 dilutions in triplicate and incubated with Lu-177 RLT (∼6.5 x 10^5^ cpm) at 37°C for 2 hours. Following incubation, supernatants were collected, and both the cell pellet and supernatant were measured using a γ-counter (Hidex AMG). The percentage of binding was calculated using decay-corrected count: (pellet – blank) / (pellet + supernatant) * 100%.

For the *ex vivo* biodistribution study, RM9-PSMA-ffLuc-bearing male C57BL/6 mice were administered with a non-therapeutic dose of Lu-177 RLT (0.37 MBq) via retro-orbital intravenous (i.v.) injection. At 0.1, 0.5, 2, 4, 6, and 168 hours post-injection, animals (n = 3 per time point) were euthanized, and tumors and organs (spleen, kidneys, gastrointestinal tract, liver, and muscle) were collected and weighed. Samples and the standard injected Lu-177 RLT were allowed to decay for 48 hours prior to γ-counting, and radioactivity was measured using a γ-counter. Tissue uptake was decay-corrected and expressed as percentage of injected dose per organ (% ID/organ). Time integrated coefficients (TIAC) for tumor and organs were estimated in MATLAB (Natick) as the area under the curve of the fraction of injected activity (non-decay corrected) vs. time curve using trapezoidal, mono- and bi-exponential fits. These calculated TIACs were then imported to the sphere model of OLINDA/EXM dosimetry package, and the output of absorbed dose vs. mass data was used to create a subset of data for masses near the appropriate organ mass. A biexponential function was then fit to the dataset to estimate the organ absorbed doses in mGy/MBq.

### Expression of PSMA and PSCA in MURAL collection and public datasets

The PDX tumor tissues were processed for histological analysis as previously described (*16*). Briefly, pathological features of PDX tumors were assessed by hematoxylin and eosin (H&E, Sigma-Aldrich) staining. PSMA and PSCA protein expression were determined by immunohistochemistry (IHC) using mouse anti-human PSMA (clone 3E6, DAKO) and mouse anti-human PSCA (polyclonal, Abcam) antibodies. PSCA transcript levels were assessed by bulk RNA-seq as previously described (*16*). Briefly, the selected PDX tumor samples were sequenced, and raw reads underwent quality control using FastQC, followed by adapter trimming by Cutadapt (v1.7.1). Reads were aligned to human reference genome hg38 using HISAT2 (v2.0.4). Gene-level counts were generated using Rsubread (v1.28.1). Normalization and expression quantification were performed using EdgeR (v3.28.1).

Processed human prostate cancer scRNA-seq data were assessed through the HuPSA resource, as previously described (*17*). Transcriptomic expression of *FOLH1* (PSMA) and *PSCA* was visualized and extracted using HuPSA-MoPSA shiny application (https://pcatools.shinyapps.io/HuPSA-MoPSA/).

### Flow cytometry analysis

Tumor cells, CAR T cells, or tumor digested single-cell suspensions were resuspended in FACS buffer (HBSS without Ca^2+^, Mg^2+^, or phenol red, supplemented with 2% FBS and 1X antibiotic-antimycotic [Gibco]). Cells harvested from mouse tumors were incubated for 20 min at 4 °C with mouse Fc block (rat anti-mouse CD16/CD32, BD Bioscience) at a 1:100 dilution prior to staining. Cells were then incubated with primary antibodies for 30 min at 4 °C in the dark, with fluorochrome-conjugated antibodies including: eFluor 506, Brilliant Violet 510 (BV510), Brilliant Violet 570 (BV570), Brilliant Violet 605 (BV605), Brilliant Violet 650 (BV650), fluorescein isothiocyanate (FITC), phycoerythrin (PE), PE/Dazzle 594, PE-cyanine (PE-Cy5), peridinin chlorophyll protein complex (PerCP), Percp-eFluor710, Percp-Cy5.5, PE-Cy7, allophycocyanin (APC), Alexa Fluor 647 (AF647), red 718 (R718), APC-Cy7, and APC-eFluor780. Cell viability was assessed using 4’, 6-diamidino-2-phenylindole (DAPI, Sigma) added immediately prior to flow cytometry acquisition.

For antigen expression, antibodies used include: APC anti-human PSMA (clone REA408/LNI-17, Miltenyi Biotec/Biolegend) and anti-human PSCA (clone 1G8, UCLA), followed by PE goat anti-mouse secondary antibody (BD Biosciences). Antigen quantification was performed using Simply Cellular anti-human IgG kit (Bangs Laboratories).

For mouse CAR T cells and tumor-derived immune cells, antibodies against mouse antigens include: PD-1 (clone J43, Invitrogen), NK1.1 (clone PK136, BioLegend), CD11b (clone M1/70, BioLegend), LAG-3 (clone C9B7W, BioLegend), CD3 (clone 17A2, BioLegend/BD Biosciences), TIM-3 (clone RMT3-23, BioLegend), CD45 (clone 30-F11, BioLegend/BD Biosciences), CD44 (clone IM7, Invitrogen), CD4 (clone RM4-5, Invitrogen), 4-1BB (clone 17B5, Invitrogen), CD62L (clone MEL-14, BioLegend), CD19 (clone 1D3, BD Biosciences), CD8a (clone 53-6.7, BioLegend), CD80 (clone 16-10A1, BD Biosciences), Ly6C (clone HK1.4, BioLegend), MHC-II (I-A/I-E) (clone M5/114.15.2, BioLegend/BD Biosciences), CD163 (clone S15049I, BioLegend), CD11c (clone N418, BioLegend), F4/80 (clone BM8, BioLegend), CD103 (clone 2E7, BioLegend), PD-L1 (clone 10F.9G2, BioLegend), Ly6G (clone 1A8, BioLegend), CD86 (clone A17199A, BioLegend), MHC-I (H-2Kb) (clone AF6-88.5, BD Biosciences), and CD206 (clone C068C2, BioLegend).

Unless otherwise specified, antibodies were diluted 1:100 for staining. Flow cytometry was performed using a MACSQuant Analyzer 16 (Miltenyi Biotec), and data were analyzed using FlowJo software (FlowJo LLC, v10.10).

### Metastatic xenograft animal model

Male NSG mice (5-6 weeks old; Jackson Laboratory) were housed in a pathogen-free facility at the University of California Los Angeles (UCLA) under a 12-hour light/dark cycle with ad libitum access to food and water. All animal studies were approved by the UCLA Institutional Animal Care and Use Committee (IACUC) and conducted in accordance with Animal Research: Reporting of In Vivo Experiments (ARRIVE) guidelines. For intracardiac injection, mice were anesthetized with isoflurane, and tumor cells were injected under ultrasound guidance. Each mouse received 0.5 × 10^6^ cells (viability >90%) in 100 μL of 1X PBS. Cells were confirmed to be mycoplasma-negative prior to injection (MycoAlert® Mycoplasma Detection Kit, Lonza). On day 15 post-inoculation, mice were treated intravenously (tail i.v.) with RLT, either [^177^Lu]Lu-PSMA-617 (Lu-177) RLT or [^225^Ac]Ac-PSMA-617 (Ac-225) RLT. PSCA-CAR T cells (0.2 × 10^6^ cells per mouse) were i.v. administered 4 or 6 days following RLT. Prior to initiation of RLT treatment and during the study, PSMA expression in vivo was evaluated with ^68^Ga-PSMA-11 PET/CT imaging.

Metastatic progression was monitored at least once weekly using an in vivo imaging system (IVIS; PerkinElmer). Bioluminescence imaging (BLI) signals were quantified using Aura in vivo imaging software (Spectral Instruments Imaging), with a consistent region of interest (ROI) applied across all time points for each animal. Mice received intraperitoneal injections of D-luciferin potassium salt (100 μL, 25 mg/mL in PBS; GoldBio) under isoflurane anesthesia 10–15 minutes prior to imaging. Hematologic toxicity was assessed by complete blood count using an Abaxis VetScan HM5 hematology analyzer (Allied Analytic). Blood samples (50 μL) were collected via retro-orbital bleeding into EDTA-coated tubes at weekly intervals up to 27 days post-RLT.

### Subcutaneous syngeneic animal model

All animal experiments were performed under protocols approved by the University of Southern California (USC) and City of Hope (COH) IACUC (protocol no. 21666 and 21687, USC; no. 21025, COH). Studies used 6-8 week-old male WT C57BL/6J (Jackson Laboratories) or heterozygous hPSCA-KI C57BL/6J mice. Mice were co-housed in a pathogen-free facility under a 12-hour light/dark cycle at 20–24°C and 30–70% humidity with ad libitum access to food and water.

For subcutaneous (s.c.) tumor studies, 1.0 x 10^6^ or 1.5 x 10^6^ RM9-derived cells (or as specified in figure legends) were resuspended in HBSS and s.c. injected into the left flank. The antigen heterogeneous tumor mixture was generated by mixing RM9-PSMA-ffLuc and RM9-PSCA-ffLuc at a 1:2 ratio prior to the tumor injection. For rechallenge studies, 0.5 x 10^6^ RM9-WT tumors were prepared in 100 µL of HBSS and s.c. injected into age-paired, tumor-naïve, or previously treated mice.

Tumor growth was monitored at least twice weekly using caliper measurements, and tumor volume was calculated using the formula: Volume = length x width x height mm^3^. Measurements continued until scheduled harvest, humane endpoints, or complete tumor regression. Where indicated, mice were treated with RLT (Lu-177 or Ac-225) via retro-orbital i.v. injection, cyclophosphamide (Cy) by i.p. administered 3 days post-RLT, and CAR T cells i.v. administered 4 days post-RLT. Mice were euthanized by CO_2_ inhalation followed by cervical dislocation upon reaching endpoints (tumor volume >1000mm^3^ or ulceration, or signs of distress including weight loss and impaired mobility).

Peripheral blood was collected as previously described (*35*). The total volume per draw (∼120 μL per mouse), or endpoint collection (∼300 μL per mouse) was recorded. CBC and serum chemistry analyses were performed as described above. Serum chemistry for kidney and liver function was analyzed using a VetScan VS2 Chemistry Analyzer (Allied Analytic) as previously described (*39*). At predetermined time points or endpoints, tumors and organs were harvested and processed for flow cytometry and IHC.

For the mechanistic studies, mice from each group were randomly selected and euthanized at day 6 post RLT injection (or as specified in the figure legends). The s.c. tumor masses were physically minced and enzymatically digested using the mouse tumor digestion kit (Miltenyi). Cells were resuspended in FACS buffer prior to flow cytometry staining or scRNA-seq processing. For the RLT + Cy + PSCA-CAR study, CD45 positive selection was performed using magnetic mouse CD45 microBeads (Miltenyi) according to manufacturer protocols. For H&E and IHC stainings, tumor or organ tissues were fixed for up to 72 hours in 4% paraformaldehyde (PFA, Boston BioProducts) and stored in 70% ethanol until further processing.

### scRNA-seq analysis

Tumors were harvested and processed into single-cell suspensions as described above. Single-cell RNA sequencing (scRNA-seq) libraries were generated using a 3′ gene expression platform (10x Genomics Chromium Single Cell kit) according to the manufacturer’s instructions. Sequencing was performed on the Novaseq X platform (Illumina) with a depth of 30,000 reads per cell. Raw data were processed using the Cell Ranger pipeline (10x Genomics) for read alignment, barcode assignment, and gene counting. Downstream analyses were conducted using R (v4.5.2). Cells with fewer than 200 or over 8000 identified genes, more than 20% of reads mapped to mitochondrial genes, were filtered out. Data were normalized and scaled using standard Seurat workflows (v5.5.0)(*40*), and dimensionality reduction was performed using principal component analysis (PCA) followed by Uniform Manifold Approximation and Projection (UMAP). Cell type annotation was performed using ScType, based on curated marker gene signatures. Differential gene expression (DEG) analysis between experimental groups was conducted using DESeq2 (v1.50.2) and edgeR (v4.8.2), with significance thresholds defined as adjusted p < 0.05 and |log₂ fold change| > 0.58 unless otherwise specified. Pathway enrichment analysis was performed using Gene Set Enrichment Analysis (GSEA) and Gene Ontology (GO) analysis. Results were visualized using running enrichment plots (GSEA), dot plots, and heatmaps. Cell–cell communication analysis was conducted using CellChat (v2.2.0.9001)(*41*), allowing inference of ligand–receptor interactions and signaling pathway activity across cell populations.

### Statistical analysis and reproducibility

All *in vitro* assays were performed with at least technical duplicate samples and were repeated in at least two independent experiments. *In vivo* studies were performed with at least three mice per group for all *in vivo* studies and downstream analyses to ensure statistical power. Mice were randomized based on bioluminescence imaging or tumour volume to ensure evenly distributed average tumour burden across each treatment group. Excel (Microsoft, v16.105.3), GraphPad Prism 11 (GraphPad Software), or R Studio (v2026.01.0) were used to generate bar plots and graphs. Data are presented as mean ± SEM. unless otherwise stated. Statistical analyses were performed using appropriate tests, including unpaired two-tailed Student’s t-tests and one- or two-way ANOVA with multiple comparison corrections as specified in the corresponding figure legends.

## List of Supplementary Materials

Figure S1-S10

Table S1

References (1–41)

## Acknowledgements

We thank the staff of the following cores at the Keck School of Medicine of the University of Southern California (USC) and the Norris Comprehensive Cancer Center: Animal Facility, EH&S Radiation Safety staff, Genomic Core and Translational Pathology Core, and the Beckman Research Institute at City of Hope (COH) Comprehensive Cancer Center: Radiopharmacy staff, Animal Facility, Pathology, and Small Animal Imaging for excellent technical assistance.

## Author contributions

S.J.P., C.E.M., and J.L. provided the conception and construction of the study. J.L., I.F., Y.R., K.P., S.Y., Y-H.F., C.A.Y., L.S.L., R.C.A.R., H.H., J.H., D.C., L.C., J.P.M, Y.Y., L.H.P., V.A., C.E.M, and S.J.P. provided the design of experimental procedures, performed experiments, data analysis, and/or interpretation. J.L. and S.J.P. wrote the manuscript. I.F., R.C.A.R., J.P.M., Y.Y., R.R., S.J.F., Y.R.L., T.B.D., G.R.R., R.T., and C.E.M. assisted in writing, editing, and reviewing the manuscript. S.J.P. supervised the study. All authors reviewed the manuscript.

## Data availability

The main data supporting the results in the study are available within the paper and its Supplementary Information.

