## Supplementary material for "Radioligand therapy in combination with CAR T cells overcomes the heterogeneous immunosuppressive prostate tumor microenvironment": Figures S1-S10, Table S1

Figure S1

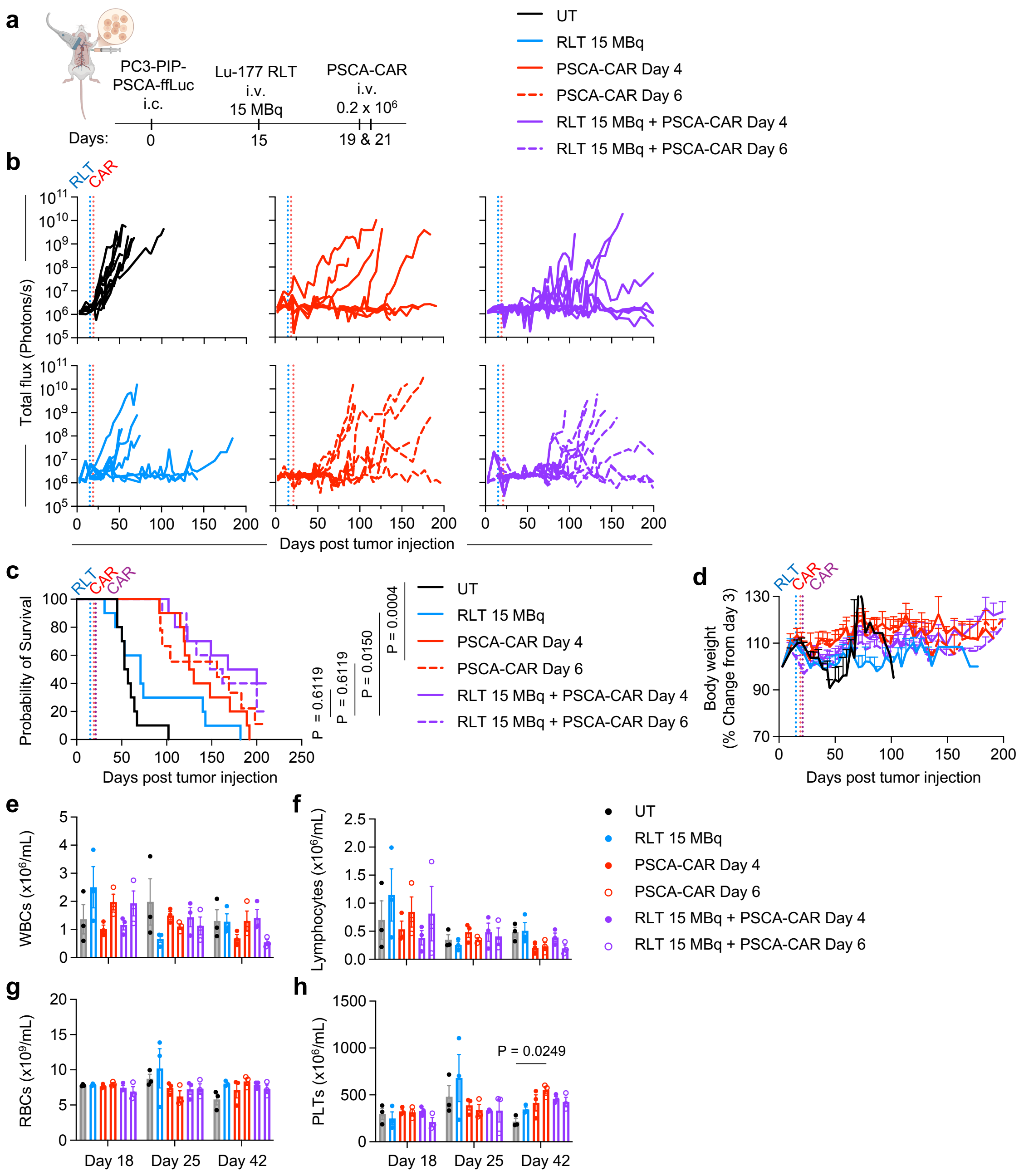


Figure S1. Modest variation in treatment intervals between Lu-177 RLT and PSCA-CAR T cell therapy does not alter therapeutic efficacy or safety in human xenograft models. (**a**) Schematic of tumor injection and treatment schedule in NSG mice bearing metastatic prostate cancer. On day 15 post-tumor injection, mice received Lu-177 RLT at 15 MBq (i.v.); Four days or six days later (day 19 or 21), mice were treated with 0.2 × $10^6$ PSCA-CAR T cells (i.v.), either as monotherapy or in combination. Blood samples (50 μL) were collected on days 18, 25, and 42 (corresponding to days 3, 7, and 21 post-RLT) for complete blood count (CBC) analysis. (**b**) Longitudinal whole-body BLI flux of PC3-PIP-PSCA-ffLuc metastatic tumor-bearing mice treated with RLT alone, PSCA-CAR T cells, or the combination (n = 10 per group). BLI was performed twice a week until day 200, and signals were quantified using a consistent region of interest (ROI) for each animal. (**c**) Kaplan-Meier survival curves for each treatment group. P-value is calculated using multiple comparisons with Holm-Šídák's correction. (**d**) Body weight changes over time in each group, normalized to body weight at day 3 post-tumor injection. Data are presented as mean ± SEM. (**e-h**) Hematologic analysis by CBC, including white blood cells (WBCs, **e**), lymphocytes (**f**), red blood cells (RBCs, **g**), and platelets (PLTs, **h**) from each group (n = 3 per group). Statistical significance was assessed using two-way ANOVA with Geisser-Greenhouse correction, followed by Tukey's multiple comparison test.

Figure S2

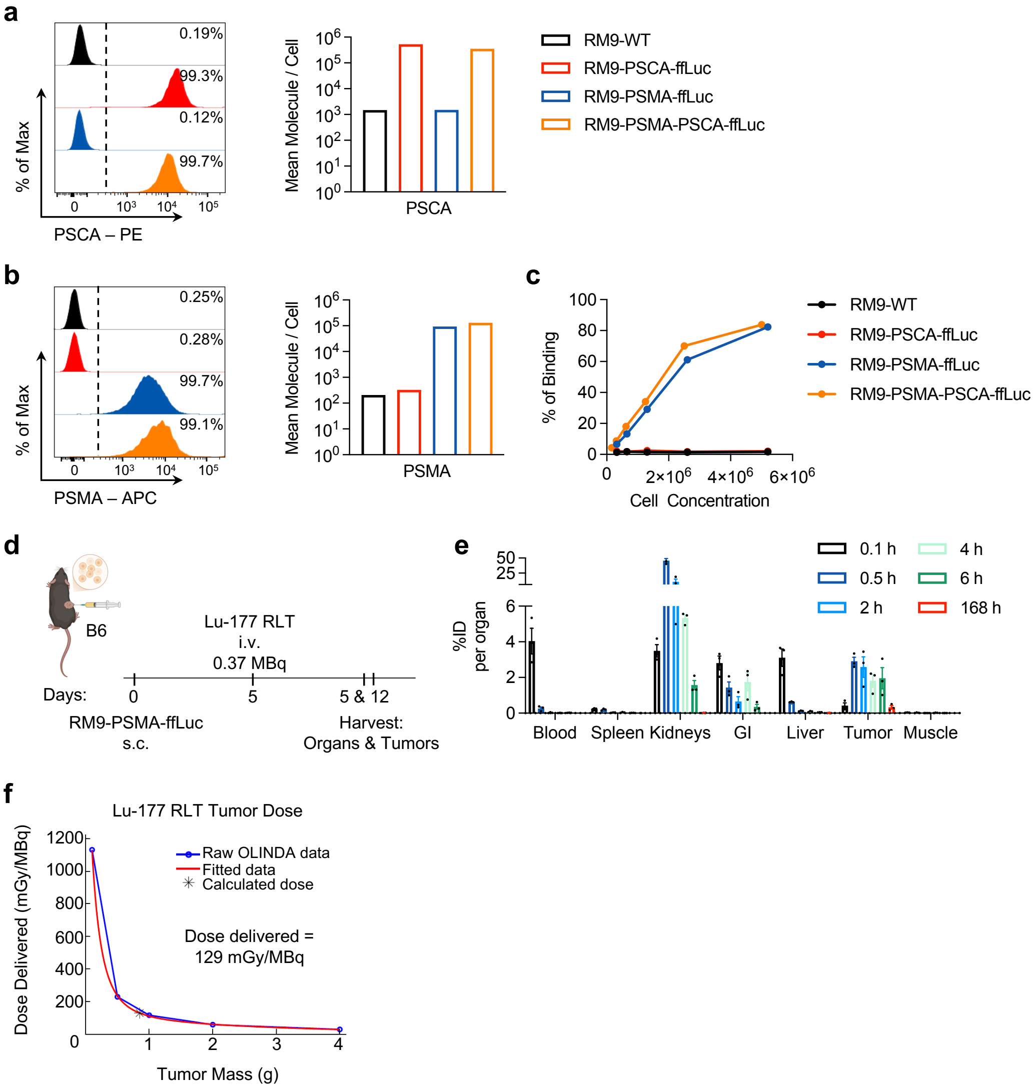


<u>Figure S2. RM9-derived model antigen expression, Lu-177 RLT binding specificity, and *in vivo* tumor targeting</u>. (**a-b**) Flow cytometry analysis of hPSCA (a) and hPSMA (b) expression in RM9-WT, RM9-PSMA-ffLuc, RM9-PSCA-ffLuc, and RM9-PSMA-PSCA-ffLuc cells, with quantification as mean molecule per cell (**c**) Binding of Lu-177 RLT to RM9 cell lines, calculated as (pellet – blank) / (pellet + supernatant) × 100%. (**d**) Schematic of biodistribution study in C57BL/6 mice bearing s.c. prostate cancer. Mice received Lu-177 RLT (0.37 KBq, i.v.) on day 5 were euthanized at 0.1, 0.5, 2, 4, 6, and 168 hours post-injection for tissue collection (n = 3 per time point). (**e**) Biodistribution of Lu-177 RLT in RM9-PSMA-ffLuc tumor-bearing mice. Data are presented as percent injected dose per organ (% ID/organ, mean ± SEM, n = 3 per time point). (**f**) Estimated

tumor dose of Lu-177 RLT in RM9-derived models. Data were computed using OLINDA/EXM from %ID/organ, presented as the mean (n = 3 per time point).

Figure S3

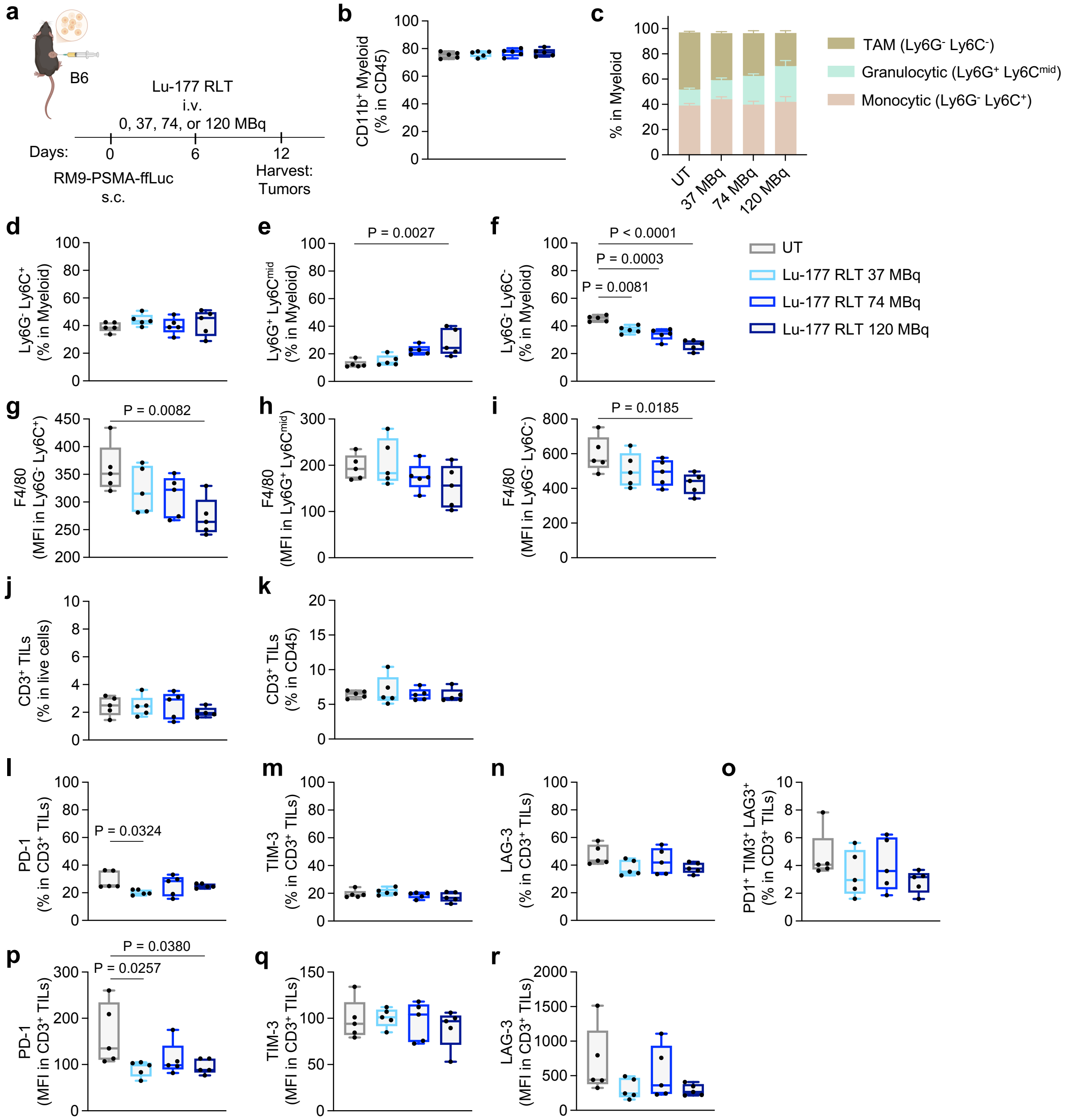


Figure S3. Lu-177 RLT modifies myeloid cell composition and endogenous T cell phenotypes in mouse syngeneic prostate cancer model. (**a**) Schematic of tumor injection and treatment schedule in C57BL/6 mice bearing subcutaneous murine prostate cancer. On day 6 post-tumor injection, mice received Lu-177 RLT (37, 74, or 120 MBq, i.v.). Tumors were harvested 6 days later (day 12) for flow cytometry (n = 5 per group). (**b-r**) Flow cytometry analysis of tumor-derived single-cell suspensions collected at 6 days post-RLT treatment (n = 5). Percent of $CD11b^+$ myeloid cells in $CD45^+$ immune cells; Composition of myeloid populations (gated on $CD11b^+$) include $Ly6G^-Ly6C^-$

tumor-associated macrophages (TAM), Ly6G$^+$Ly6C$^{mid}$ granulocytic cells, and Ly6G$^-$Ly6C$^+$ monocytic cells shown as summary (**c**) and individually (**d-f**); mean fluorescence intensity (MFI) of F4/80 in individual myeloid populations (**g-i**); percentage of CD3$^+$ in total live cells (**j**) and CD45$^+$ immune cells (**k**); percentage of checkpoint markers PD-1 (**l**), TIM-3 (**m**), LAG-3 (**n**), and PD-1$^+$ TIM-3$^+$ LAG-3$^+$ triple positive T cells (**o**); and mean fluorescence intensity (MFI) of PD-1 (**p**), TIM-3 (**q**), and LAG-3 (**r**) in T cells. Box plots represent median with interquartile range and whiskers indicating minimum and maximum; bar graphs represent mean ± SEM. Statistical significance was determined by one-way ANOVA with Bonferroni correction.

Figure S4

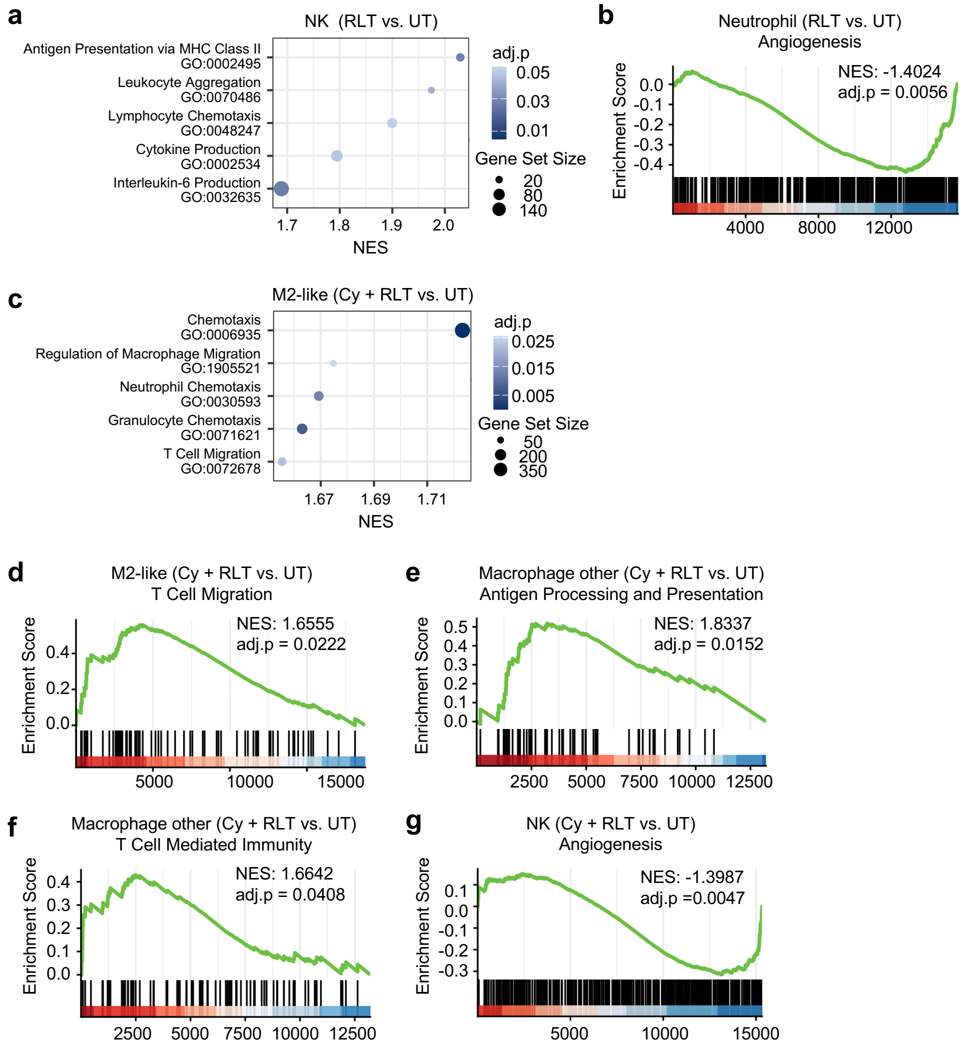


Figure S4. Lu-177 RLT alone or with Cy induces broad transcriptional remodeling of myeloid cell populations. (**a**) Representative top GSEA results for NK cells comparing RLT versus untreated (UT) groups, ranked by normalized enrichment score (NES). (**b**) Representative GSEA enrichment plot for pathway enrichment in Neutrophils from RLT versus UT. (**c**) Representative top GSEA results for M2-like macrophages comparing Cy + RLT versus UT, ranked by normalized enrichment score (NES). (**d-g**) Representative GSEA enrichment plot for pathway enrichment in M2-like macrophages (**d**), macrophages - other (**e-f**), and NK cells (**g**) from Cy + RLT versus UT.

Figure S5

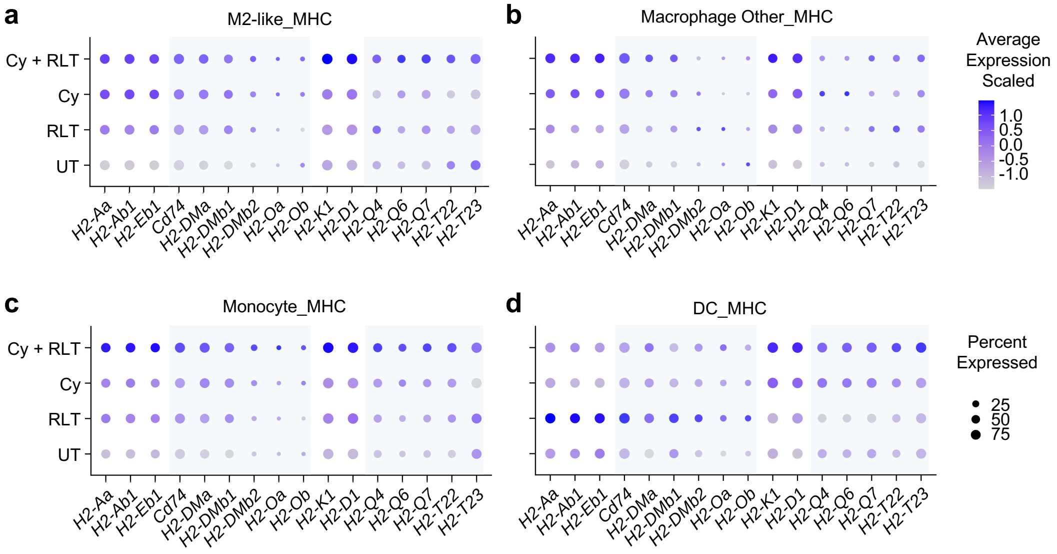


Figure S5. Lu-177 RLT alone or with Cy enhances antigen presentation programs across myeloid cell populations. (**a-d**) Balloon plot of MHC class I and class II genes (classical and non-classical) in M2-like macrophages (**a**), macrophages – other (**b**), monocytes (**c**), and dendritic cells (DCs, **d**) across treatment groups (UT, RLT, Cy, and Cy + RLT). Dot size represents the proportion of expressing cells, and color intensity reflects relative expression.

Figure S6

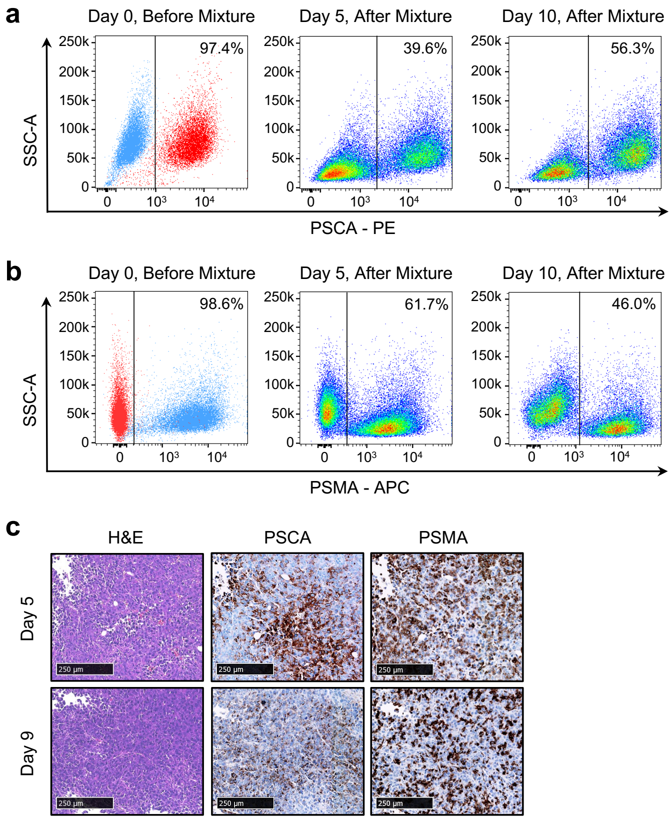


<u>Figure S6. Antigen heterogeneity is maintained in mixed RM9 models *in vitro* and *in vivo*</u>. (**a-b**) Flow cytometry analysis of hPSCA (**a**) and hPSMA (**b**) expression in mixed RM-9 cell populations. RM9-PSMA-ffLuc (blue) with RM9-PSCA-ffLuc (red) cells were combined to generate a heterogeneous population and analyzed at baseline (day 0, prior to mixing) and after 5 and 10 days of co-culture. (**c**) Representative H&E and Immunohistochemistry (IHC) staining of heterogeneous tumors collected *in vivo* at day 5 (RLT treatment) and day 9 (CAR T treatment), showing for hPSCA and hPSMA. The tumors were collected from tumor-bearing untreated mice (n = 3).

Figure S7

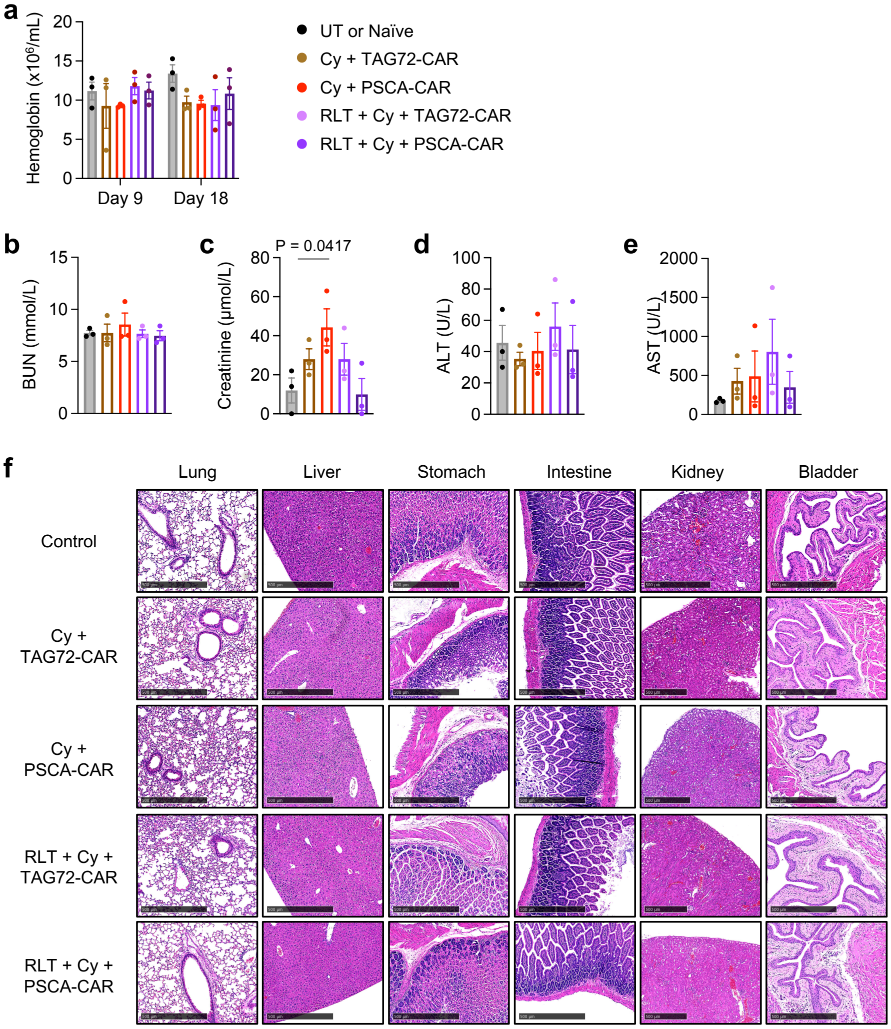


<u>Figure S7. Combination therapy demonstrates a favorable safety profile</u>. (**a**) Hematologic analysis by CBC of hemoglobin (n = 3 per group). Tumor-bearing mice received Lu-177 RLT (74 MBq, i.v.) on day 5, followed by Cy (100 mg/kg, i.p.) on day 8, and CAR T cells (PSCA-CAR or TAG72-CAR control, 1 × $10^6$, i.v.) on day 9. Blood samples were collected on day 9 (prior to CAR T cell treatment) and day 18 (tumor regression phase). Controls included tumor-bearing untreated (UT) mice at day 9, and tumor-naïve mice at day 18. Statistical significance was assessed using two-way ANOVA, followed by Tukey's multiple comparison test. (**b-e**) Serum chemistry analysis, including blood urea nitrogen (BUN, **b**) and creatinine (CRE, **c**), as indicators of kidney function, and alanine aminotransferase (ALT, **d**) and aspartate aminotransferase (AST, **e**), as indicators of liver function. Statistical significance was determined by one-way ANOVA with Bonferroni correction. (**f**) Representative H&E staining of major organs, including lung, liver, stomach, intestine, kidney, and bladder, collected from mice treated with Cy + TAG72-CAR, Cy + PSCA-

CAR, RLT + Cy + PSCA-CAR, and RLT + Cy + PSCA-CAR on day 18. Age-matched healthy mice served as controls (n = 3 per group).

Figure S8

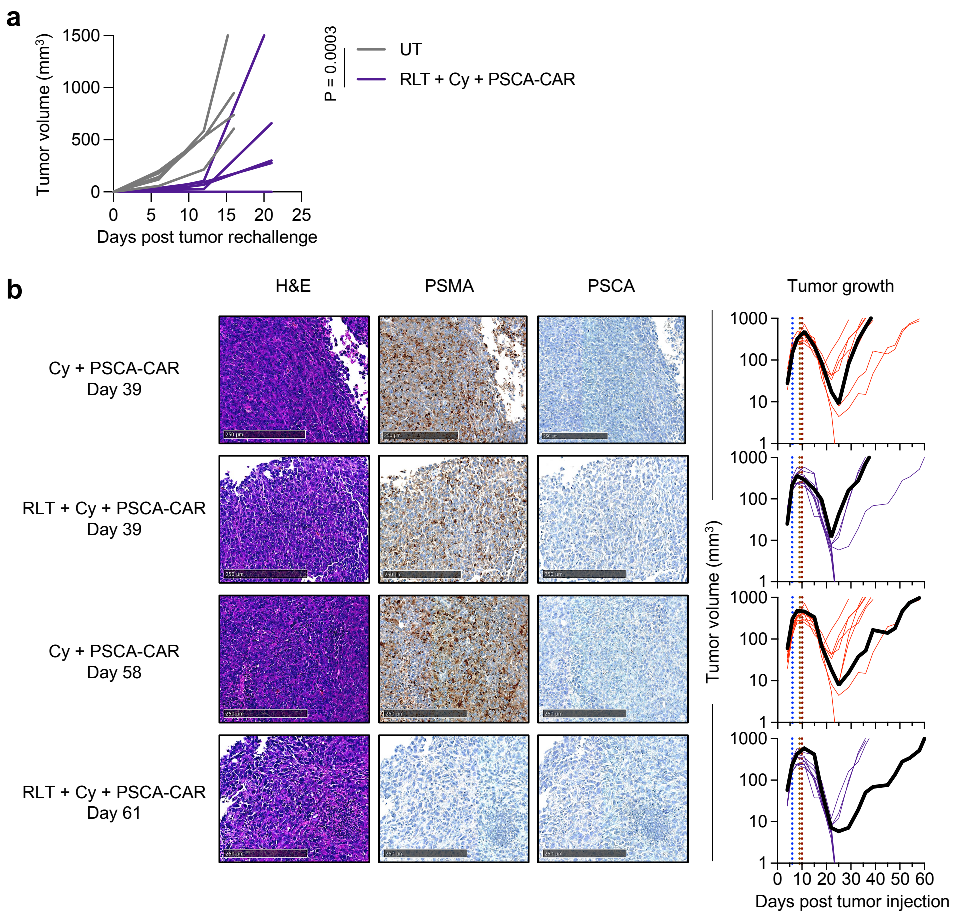


Figure S8. Combination therapy induces antigen-independent immune memory and reveals potential resistance mechanisms. (**a**) Individual tumor volume of mice rechallenged with RM9-WT (0.5 × $10^6$, s.c.). Cured mice (n = 5) previously treated with RLT + Cy + PSCA (Figure 5) were rechallenged with RM9-WT, and age-matched, treatment-naïve mice served as controls (n = 5). Statistical significance was assessed by unpaired t-test with multiple comparisons correction. P-value shown corresponds to comparisons at day 12 post tumor rechallenge. (**b**) Representative H&E and IHC staining of tumors collected at survival endpoints from mice treatment with Cy + PSCA-CAR or RLT + Cy + PSCA-CAR, including early relapse (day 39) and late relapse (day 58-61), showing hPSCA and hPSMA. The rightmost panel shows representative growth curves corresponding to each condition. At least two animals per time point (or comparable time frame) were analyzed.

Figure S9

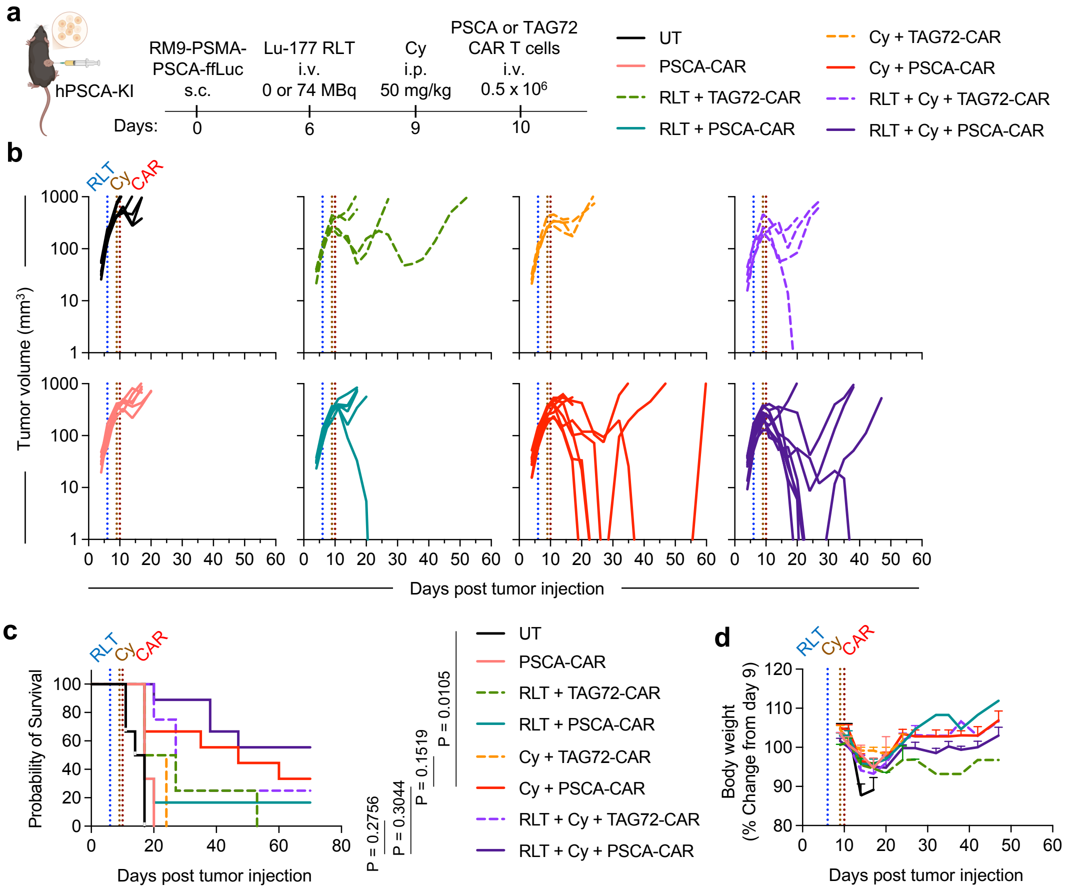


Figure S9. Combination therapy improves tumor control in homogeneous antigen-expressing syngeneic model. (**a**) Schematic of tumor injection and treatment schedule in hPSCA-KI C57BL/6 mice bearing homogeneous subcutaneous prostate tumors. 1 × $10^6$ RM9-PSMA-PSCA-ffLuc cells were engrafted into the left flank. On day 6 post-tumor injection, mice received Lu-177 RLT (74 MBq, i.v.), followed by Cy (50 mg/kg, i.p.) on day 8, and CAR T cells (PSCA-CAR or TAG72-CAR non-targeting control, 5 × $10^5$, i.v.). (**b**) Individual tumor growth curves. Tumor volume was measured by caliper twice weekly until the survival endpoint (n = 9 for RLT + Cy + PSCA-CAR/TAG72-CAR; n = 6 for RLT + PSCA-CAR and PSCA-CAR only; n = 4 for the remaining control groups). (**c**) Kaplan-Meier survival curves for each treatment group. P-value is calculated using multiple comparisons with Holm-Šídák's correction. (**d**) Body weight changes over time, normalized to baseline (day 9, prior to CAR T cell treatment). Data are presented as mean ± SEM.

Figure S10

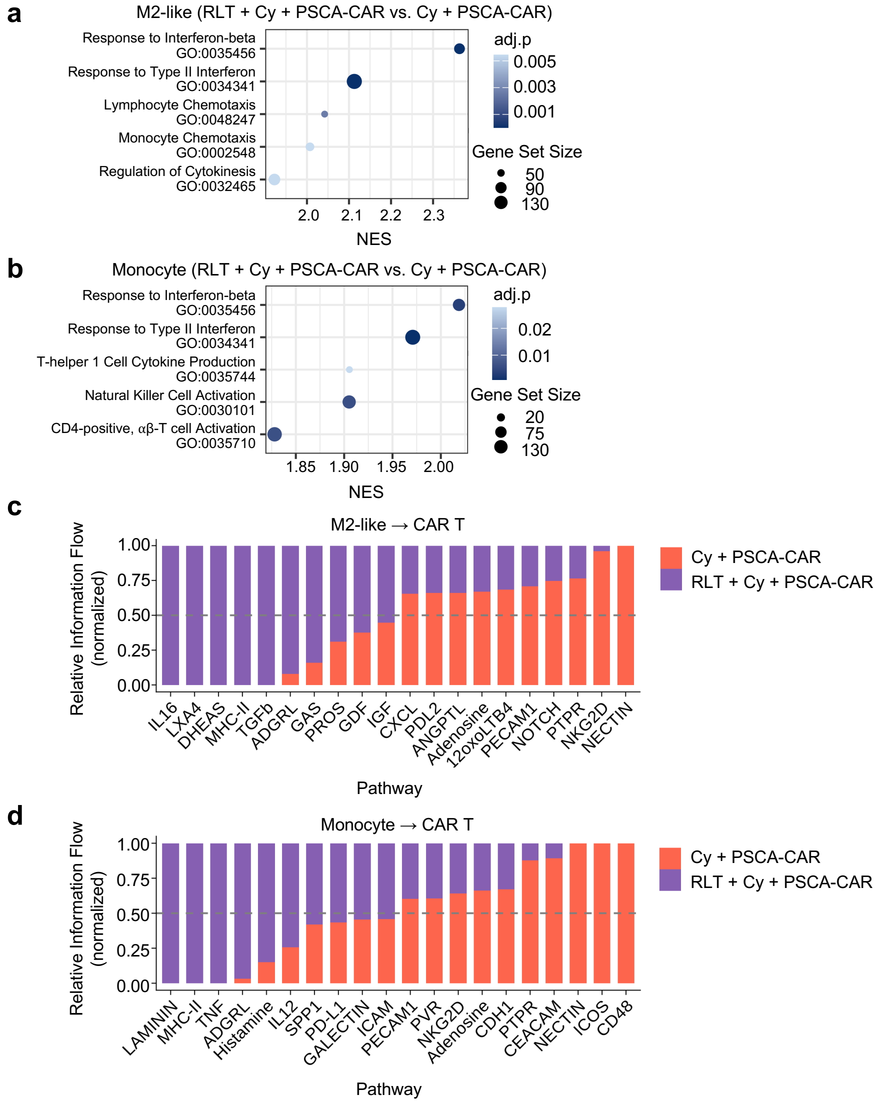


<u>Figure S10. Lu-177 RLT enhances myeloid cell-CAR T cell communication and promotes pro-inflammatory signaling networks</u>. (**a-b**) Representative top GSEA of enriched pathways in M2-like macrophages (**a**), and monocytes (**b**) comparing RLT + Cy + PSCA-CAR versus Cy + PSCA-CAR groups, ranked by normalized enrichment score (NES). (**c-d**) Cell-cell communication analysis by CellChat, showing signaling interactions from M2-like macrophages (**c**) or monocytes (**d**) as ligands, to CAR T cells as receptors. Bar plots represent relative information flow, normalized across conditions, with the top 10 pathways shown for each group.

Table S1

| Organ | Organ mass (g) | Dose (mGy/MBq) |
|---|---|---|
| Spleen | 0.2 | 1.8 |
| Kidneys | 0.5 | 124.2 |
| GI | 3.3 | 21.3 |
| Liver | 1.8 | 3.3 |
| Tumor | 0.9 | 129.4 |
| Muscle | 0.1 | 15.5 |

Table S1. Estimated dose of Lu-177 RLT in RM9-derived models. Data were computed using OLINDA/EXM from %ID/organ, presented as the mean (n = 3 per time point).
